# Genome-wide detection of human variants that disrupt intronic branchpoints

**DOI:** 10.1101/2022.04.18.488668

**Authors:** Peng Zhang, Quentin Philippot, Weicheng Ren, Wei-Te Lei, Juan Li, Peter D. Stenson, Pere Soler Palacín, Roger Colobran, Bertrand Boisson, Shen-Ying Zhang, Anne Puel, Qiang Pan-Hammarström, Qian Zhang, David N. Cooper, Laurent Abel, Jean-Laurent Casanova

## Abstract

Pre-mRNA splicing is initiated with the recognition of a single-nucleotide intronic branchpoint (BP) within a BP motif by spliceosome elements. Fifty-six rare variants in 44 human genes have been reported to alter splicing and cause disease by disrupting BP. However, until now, no computational approach has been available to efficiently detect such variants in next-generation sequencing (NGS) data. We established a comprehensive human genome-wide BP database by integrating existing BP data, and by generating new BP data from RNA-seq of lariat debranching enzyme DBR1-mutated patients and from machine-learning predictions. We in-depth characterize multiple features of BP in major and minor introns, and find that BP and BP-2 (two-nucleotides upstream of BP) positions exhibit a lower rate of variation in human populations and higher evolutionary conservation than the intronic background, whilst being comparable to the exonic background. We develop BPHunter as a genome-wide computational approach to systematically and efficiently detect intronic variants that may disrupt BP recognition in NGS data. BPHunter retrospectively identifies 48 of the 56 known pathogenic BP mutations in which we summarize a strategy for prioritizing BP mutation candidates, and the remaining 8 all create AG dinucleotides between BP and acceptor site which is probably the reason for mis-splicing. We demonstrate the utility of BPHunter prospectively by using it to identify a novel germline heterozygous BP variant of *STAT2* in a patient with critical COVID-19 pneumonia, and a novel somatic intronic 59-nucleotide deletion of *ITPKB* in a lymphoma patient, both of which we validate experimentally. BPHunter is publicly available from https://hgidsoft.rockefeller.edu/BPHunter and https://github.com/casanova-lab/BPHunter.

## INTRODUCTION

Pre-mRNA splicing is a necessary step for protein-coding gene expression in eukaryotic cells. The biochemical process is initiated by the recognition of the branchpoint (BP) in an intron, followed by the identification and ligation of the 5’ splice site (5’ss = donor site) and 3’ splice site (3’ss = acceptor site) to join two exons, and the removal of the intervening intron as a circular lariat, to yield a mature mRNA of coding sequence for protein translation (**Figure 1a**). Splicing takes place in the nucleus, and is regulated by an array of *cis*-acting elements (the RNA sequences with their splicing codes) and *trans*-acting elements (proteins and small nuclear RNAs (snRNAs) that bind to the *cis*-acting elements), which together constitute the complex and dynamic cellular spliceosome (Wang and Cooper 2007; Scotti and Swanson 2016). When splicing is initiated, the BP motif is recognized by a spliceosomal snRNA, with the cooperation of splicing factors (Gao et al. 2008; Mercer et al. 2015). BP recognition is achieved differently between the two types of spliceosome (**Figure 1b**): the major (U2-type, in >99% of human introns) and the minor (U12-type). The main difference between these spliceosomes is the use of different snRNAs. The major spliceosome recruits U1, U2, U4, U5 and U6 snRNAs, in which the U1 and U2 snRNAs recognize 5’ss and BP independently (Scotti and Swanson 2016). In the interaction between BP and U2 snRNA, the BP nucleotide bulges out to bind into a pocket formed by SF3B1 and PHF5A (Tholen et al. 2022), and its flanking sequences base-pair with U2 snRNA (Turunen et al. 2013; Mercer et al. 2015), while U2 snRNA is stabilized by SF3B6 (Tholen et al. 2022). The minor spliceosome recruits U11, U12, U4atac, U5 and U6atac snRNAs, in which the U11 and U12 snRNAs first form a di-snRNA complex before recognizing the 5’ss and BP simultaneously (Turunen et al. 2013; Bai et al. 2021).

**Figure 1:**
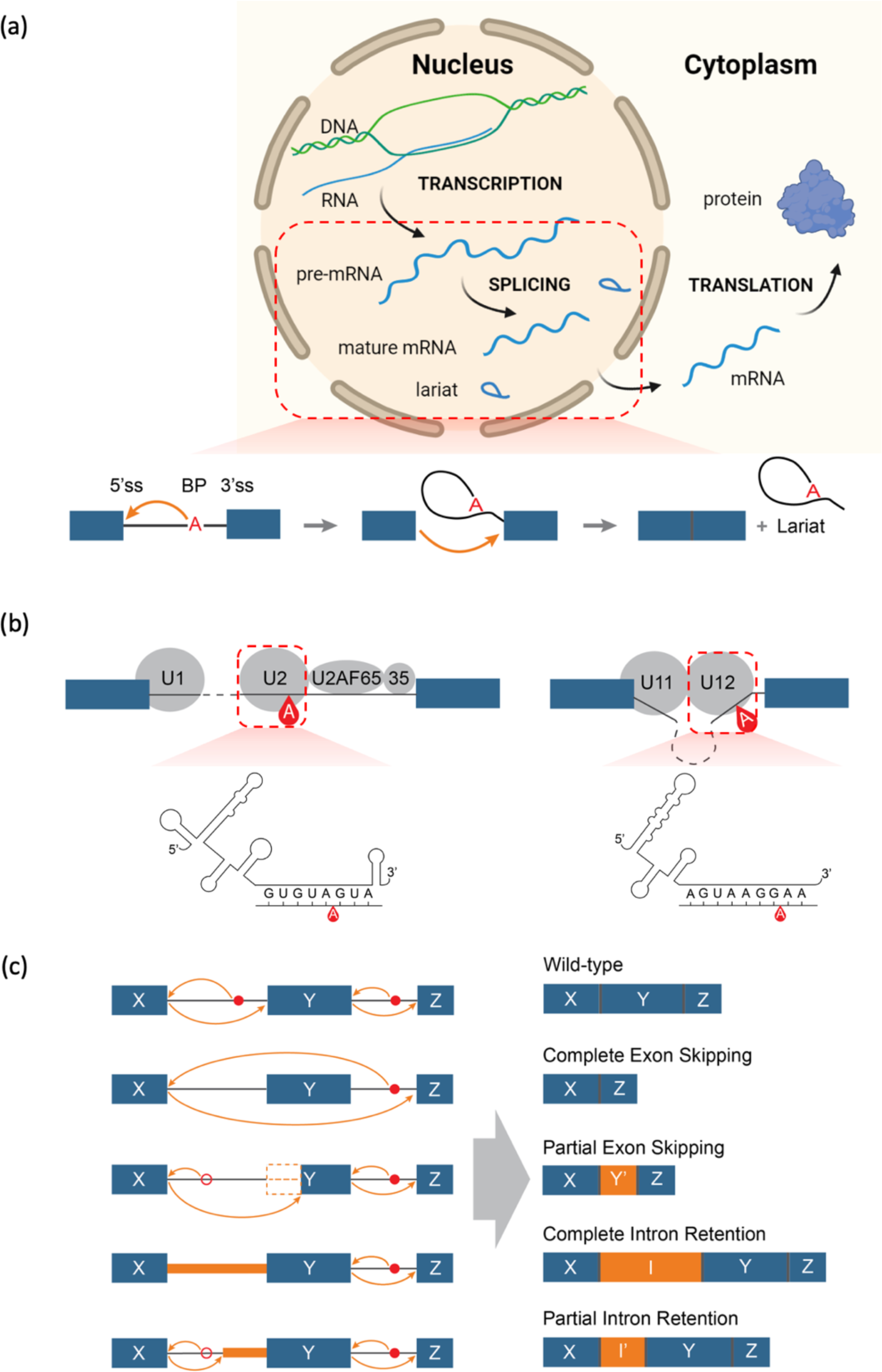
Schematics of a splicing event, the two types of spliceosome, and typical consequences of a branchpoint (BP) mutation. (a) Schematic of the biological processes of transcription, splicing and translation from DNA to pre-mRNA to mature mRNA and protein. In pre-mRNA splicing, the BP is first recognized, two exons are then joined, and the intervening intron is finally released as a circular lariat. (b) Schematic of the major (U2-type, on the left) and minor (U12-type, on the right) spliceosomes, with an illustration of the interaction between pre-mRNA sequence and U2/U12 snRNA. (c) Schematic of the potential molecular consequences of a BP mutation, including complete/partial exon skipping and complete/partial intron retention.

BP constitutes a single-nucleotide *cis*-acting element within the intron. It is typically an adenine and generally located [-20, -40] (from -20 to -40) nucleotides (nt) away from the 3’ss. The BP motif is thought to follow the consensus sequence YTN<u>A</u>Y (Gao et al. 2008) (<u>A</u> denotes the BP site, Y = C/T, N = any nucleotide). There are also distal BP deeper inside introns: a recursive splicing mechanism could use multiple intermediate 5’ss-BP-3’ss in an intron for multi-step intron removal (Sibley et al. 2015; Wan et al. 2021); stem-loop RNA structures could bring distal BP closer to 3’ss to facilitate splicing (Lin et al. 2016) (**Figure S1**). Considerable efforts have been made to identify BP positions, including developing a lariat sequencing method to study BP in yeast (Awan et al. 2013), analyzing large-scale experimental data in human (RNA-seq (Taggart et al. 2012; Mercer et al. 2015; Taggart et al. 2017; Pineda and Bradley 2018; Talhouarne and Gall 2018) and iCLIP (Briese et al. 2019)), and making human genome-wide computational predictions (Zhang et al. 2017; Paggi and Bejerano 2018; Signal et al. 2018). To locate BP positions from RNA-seq data, the sequencing reads from the circular lariats that traverse the junctions between 5’ss and BP (5’ss-BP junction reads) are identified by aligning the reads to the intronic sequences of 5’ss and 3’ss respectively. Spliceosome iCLIP experiment identifies crosslinks of spliceosomal factors on pre-mRNA sequences at nucleotide-level resolution to pinpoint BP positions (Briese et al. 2019). Some machine-learning models have also been developed based on one or two experimentally identified BP datasets, and have been used to predict BP positions in regions upstream of 3’ss. However, these individual BP studies have yet to be integrated into a comprehensive database and knowledgebase of BP.

The functional role of splicing makes it a critical process for the correct assembly of gene products, whilst its complexity renders it vulnerable to deleterious mutations that can perturb any part of the splicing machinery. A mutation may affect a *cis*- or *trans*-element, leading to altered splicing with alternative 5’ss/3’ss, exon skipping or intron retention, any of which could result in the synthesis of a defective gene product (Cooper et al. 2009; Singh and Cooper 2012). As previously reported (Sterne-Weiler et al. 2011; Bao et al. 2019; Stenson et al. 2020), at least 10% of disease-causing mutations are associated with mis-splicing (of which ∼65% are in essential exonic/intronic splice sites, ∼25% in the remainder of the intron, and ∼10% in the remainder of the exon (Stenson et al. 2020)); this may however be a serious underestimate (Bao et al. 2019) owing to our rather limited ability to detect mutations that impair different aspects of spliceosome. Currently, the search for candidate pathogenic mutations in next-generation sequencing (NGS) data generally focuses on non-synonymous variants located within coding sequences or variants in essential splice sites, while mostly ignoring synonymous coding exonic variants and non-coding intronic variants. Data from RNA-seq or other studies of mRNA occasionally lead to the detection of intronic or exonic variants that disrupt splicing (Kremer et al. 2017). However, the spliceosome operates mostly within intronic regions to define the exon-intron boundaries and hence the coding sequences. It follows that introns probably harbor a substantially larger number of pathogenic mutations than has so far been appreciated (Cooper 2010). The BP, involved in the initiation of splicing through its interactions with spliceosome elements, constitutes a key vulnerability of splicing by virtue of its potential mutations (Kralovicova et al. 2006).

Employing the Human Gene Mutation Database Professional v.2021.2. (HGMD) (Stenson et al. 2020) in concert with an in-depth literature review, we identified 56 BP mutations reported in 44 genes underlying 41 human disease whose pathogenicity has been supported by accompanying experimental evidence. These BP mutations resulted in complete/partial exon skipping or intron retention, or a combination of these mis-splicing consequences (**Figure 1c**). The mis-spliced mRNAs may be rapidly degraded by nonsense-mediated decay (NMD) resulting in loss of expression, and the translated truncated proteins could exhibit loss of function, both being potentially disease-causing. However, the discovery rate of pathogenic BP mutations (only 56 mutations discovered in the last 30 years) appears to have been rather low. There were some studies attempted to use different BP datasets to identify and test BP mutations in cancer genes (Leman et al. 2020; Canson et al. 2021), but we still lack some important information, tools and guides to support the investigation of BP mutations in human disease: (1) genome-wide knowledge of BP has not been systematically integrated and characterized; (2) genome-wide detection software has not been available for an efficient identification of BP variants from NGS data; and (3) in-depth understanding of all reported pathogenic BP mutations has not been analyzed, which could provide strategic prioritization of variant candidates that could functionally disrupt BP. We therefore have aimed to fill these gaps both for biological and medical reasons.

## RESULTS

### Collection of experimentally identified BP (eBP) data

We collated five datasets of experimentally identified BP, based on RNA-seq and iCLIP data: eBP_Mercer (59,359 BP from total RNA-seq) (Mercer et al. 2015), eBP_Taggart (27,795 BP from total RNA-seq) (Taggart et al. 2017), eBP_Pineda (138,313 BP from total RNA-seq) (Pineda and Bradley 2018), eBP_Talhouarne (240 BP from cytoplasmic RNA-seq) (Talhouarne and Gall 2018) and eBP_Briese (43,630 BP from iCLIP) (Briese et al. 2019) (**Table 1**). From RNA-seq data, BP were identified based on the reads from lariats that traverse the 5’ss-BP junctions. Since lariats are transient and only exist at low abundance, their detection is technically difficult, hence large-scale data are required to profile the genome-wide distribution of BP (Pineda and Bradley 2018). In addition to the published BP datasets, we revisited the RNA-seq data derived from *DBR1*-mutated patients with brainstem viral infection, which our group previously studied (Zhang et al. 2018b). *DBR1* encodes the only known lariat debranching enzyme, and the patients with *DBR1* mutations, who were characterized by a residual DBR1 activity of 3-10%, were reported to exhibit an elevated cellular level of lariats (Zhang et al. 2018b). We designed a computational program to detect BP positions from RNA-seq reads (**Methods**), which identified 8,682 BP (eBP_BPHunter) from 15 RNA-seq datasets from the fibroblasts of three *DBR1*-mutated patients (**Figure 2a, Table S1**). In these six eBP datasets, the most abundant BP nucleotides were: eBP_Mercer (78% A), eBP_Taggart (68% A), eBP_Pineda (75% A), eBP_Talhouarne (84% C), eBP_Briese (100% A) and eBP_BPHunter (46% A) respectively (**Table 1**). This showed that adenine-BP were the most frequent in lariats from total RNA-seq, whereas cytosine-BP were the most frequent in the stable lariats in cytoplasm albeit in very low amounts (Talhouarne and Gall 2018). About one half of BP from *DBR1*-mutated patients were adenine whilst having 22% stable cytosine-BP. The iCLIP-identified BP were invariably adenine. In total, we obtained 210,986 unique eBP.

**Figure 2:**
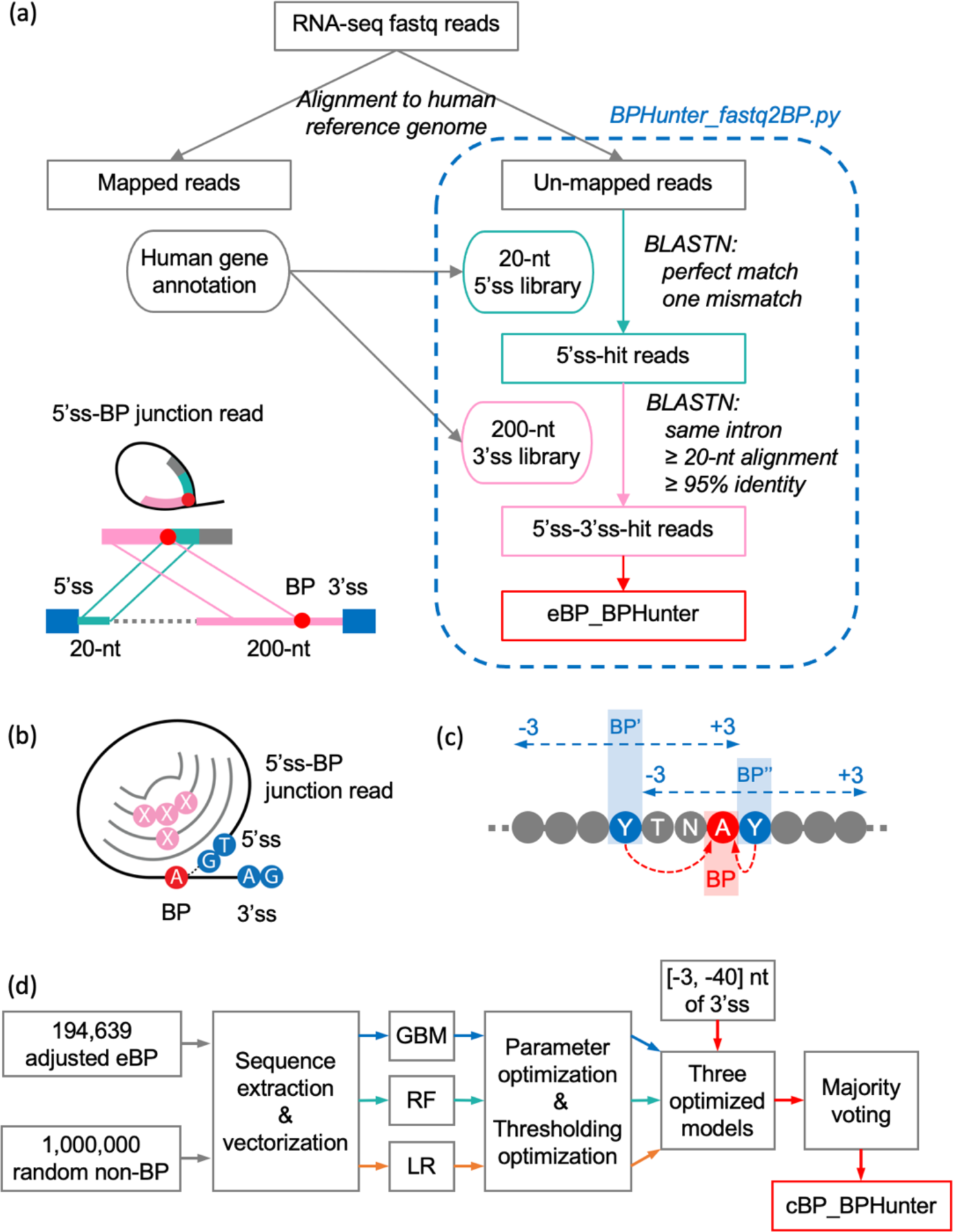
Identification of eBP_BPHunter, BP positional adjustment, and prediction of cBP_BPHunter. (a) Workflow of BP identification from RNA-seq reads from *DBR1*-mutated patients. (b) Introduction of mutations to the 5’ss-BP junction reads by reverse transcriptase in an RNA-seq experiment. (c) Positional adjustment of BP within its [-3, +3] neighborhood, guided by the BP consensus sequence (blue: raw BP position, red: adjusted BP position). (d) Development of three machine learning models to majority-voted prediction of BP within the region [-3, -40] nt of 3’ss.

**Table 1:** A comprehensive collection of experimentally identified BP (eBP) and computationally predicted BP (cBP), including the additional BP data generated in this study (eBP_BPHunter and cBP_BPHunter), followed by consensus-guided positional adjustment, for a total of 567,053 BP positions, with 104,349 mutually shared (mBP) between eBP and cBP.

| BP dataset | Raw BP |  |  |  |  | Adjusted BP |  |  |  |  |  |
| --- | --- | --- | --- | --- | --- | --- | --- | --- | --- | --- | --- |
| | # BP | A | C | G | T | # BP | A | C | G | T | $\Delta$ #BP |
| eBP_Mercer | 59,359 | 78.45% | 8.45% | 4.73% | 8.38% | 53,784 | 85.60% | 7.65% | 3.30% | 3.44% | -5,575 |
| eBP_Taggart | 27,795 | 67.82% | 15.29% | 5.54% | 11.35% | 27,450 | 76.18% | 12.24% | 3.91% | 7.67% | -345 |
| eBP_Pineda | 138,313 | 75.02% | 7.57% | 9.52% | 7.89% | 131,676 | 77.86% | 6.84% | 8.85% | 6.46% | -6,637 |
| eBP_Talhouarne | 240 | 4.58% | 84.17% | 4.58% | 6.67% | 240 | 12.50% | 77.92% | 3.33% | 6.25% | 0 |
| eBP_Briese | 43,630 | 100% | 0% | 0% | 0% | 43,630 | 100% | 0% | 0% | 0% | 0 |
| eBP_BPHunter | 8,682 | 45.93% | 22.39% | 18.20% | 13.48% | 7,829 | 59.75% | 21.19% | 13.44% | 5.62% | -853 |
| **eBP** | 210,986 | 73.39% | 9.06% | 8.49% | 9.06% | 194,639 | 77.99% | 8.15% | 7.52% | 6.35% | -16,347 |
| cBP_BPP | 232,828 | 98.76% | 1.05% | 0.00% | 0.19% | 232,819 | 98.79% | 1.03% | 0.00% | 0.17% | -9 |
| cBP_Branchpointer | 352,322 | 98.81% | 0.50% | 0.01% | 0.67% | 320,016 | 99.45% | 0.55% | 0.00% | 0.00% | -32,306 |
| cBP_LaBranchoR | 203,914 | 98.31% | 1.63% | 0.03% | 0.03% | 203,903 | 98.35% | 1.60% | 0.03% | 0.02% | -11 |
| cBP_BPHunter | 105,808 | 100% | 0% | 0% | 0% | 105,808 | 100% | 0% | 0% | 0% | 0 |
| **cBP** | 512,414 | 98.04% | 1.38% | 0.02% | 0.56% | 476,763 | 98.44% | 1.46% | 0.01% | 0.09% | -35,651 |
| **Total BP** |  |  |  |  |  | 567,053 | 91.28% | 3.87% | 2.59% | 2.26% |  |
| **mBP** |  |  |  |  |  | 104,349 | 99.17% | 0.81% | 0.01% | 0.01% |  |

### Collection of computationally predicted BP (cBP) data

We also collated three datasets of computationally predicted BP, based on machine-learning models trained on one or two eBP datasets, and performed predictions in the region [-21, -34], [-18, -44] and [-1, - 70] nt of 3’ss respectively: cBP_BPP (232,828 BP) (Zhang et al. 2017), cBP_Branchpointer (352,322 BP) (Signal et al. 2018) and cBP_LaBranchoR (203,914 BP) (Paggi and Bejerano 2018) (**Table 1**), which supplemented the BP positions that were not detected from experimental data and hence increased the BP coverage across the human genome. These computational methods overlapped the predictions in [-21, -34] nt of 3’ss, with less attention in the region closer to 3’ss, which is generally considered to comprise polypyrimidine tracts (PPT). However, the use of a hard distance cutoff for PPT or the use of an algorithm to estimate PPT to skip BP prediction, could miss potentially genuine BP sites. For this reason, we postulated that there could be additional BP candidates in the region closer to 3’ss. Therefore, we developed a set of three machine-learning models (gradient boost machine (GBM), random forest (RF), and logistic regression (LR)) with majority voting for the final prediction (**Methods**). These three models were independently trained on the 194,639 consensus-guided position-adjusted eBP positions (discussed in the next section, **Figure 2b** and **2c**) versus 1,000,000 random intronic/exonic positions, and then optimized by tuning parameters and thresholds through cross-validation (**Figure 2d**). We aimed to achieve high precision in optimizing each model, and used the optimized models to predict BP in [-3, -40] nt of 3’ss. As a result, our prediction yielded 105,808 BP (cBP_BPHunter). In these four cBP datasets, the most abundant BP nucleotides were: cBP_BPP (99% A), cBP_Branchpointer (99% A), cBP_LaBranchoR (98% A) and cBP_BPHunter (100% A) respectively (**Table 1**), showing that the prediction models favored adenine-BP. In total, we obtained 512,414 unique cBP. It should be noted that cBP does not denote cytosine-BP. When we mention the nucleotide of BP in this article, we fully spell out the nucleotide (e.g., adenine-BP, cytosine- BP).

### Consensus-guided positional adjustment of BP, and integration of BP datasets

We tested the eBP against the established BP consensus sequence YTNAY (Gao et al. 2008), but found only 23.5% eBP matched this pattern precisely. Considering the noncanonical 2’-to-5’ linkage between 5’ss and BP, and the possibility of mutations being introduced by reverse transcriptase when traversing the 5’ss-BP junction in RNA-seq (Mercer et al. 2015; Taggart et al. 2017), we anticipated that a number of eBP may have been mis-located in the raw dataset (**Figure 2b**), which could also have led to some mis-located cBP predictions. We therefore screened a window of [-3, +3] nt from each BP position for consensus sequence matching, and adjusted the raw BP position to its closest neighbor that perfectly matched the consensus YTNAY (**Figure 2c**). If the YTNAY pattern was not found, the slightly relaxed consensus YTNA was used for positional adjustment. As a result, 11.8% raw eBP positions and 5.7% raw cBP positions were adjusted (**Table S2**), and multiple neighboring raw BP positions were merged into a single adjusted BP position, yielding 194,639 eBP (Δ = -16,347) and 476,763 cBP (Δ = -35,651) after adjustment (**Table 1**). By integrating eBP and cBP, we assembled a comprehensive collection of 567,053 BP in the human genome, with 104,349 mutually shared BP (mBP) (**Figure 3a, Table 1, Table S3, Supplemental Data 1**). This BP database contained 91.3% adenine-BP, 3.9% cytosine-BP, 2.6% guanine- BP, and 2.3% thymine-BP. Among adenine-BP, we decomposed adenine-BP by adopting increasingly relaxed consensus sequences: YTNAY (40.6%), YTNA (52.8%), TNA (69.7%) and YNA (79.1%) (**Figure S2**).

**Figure 3:**
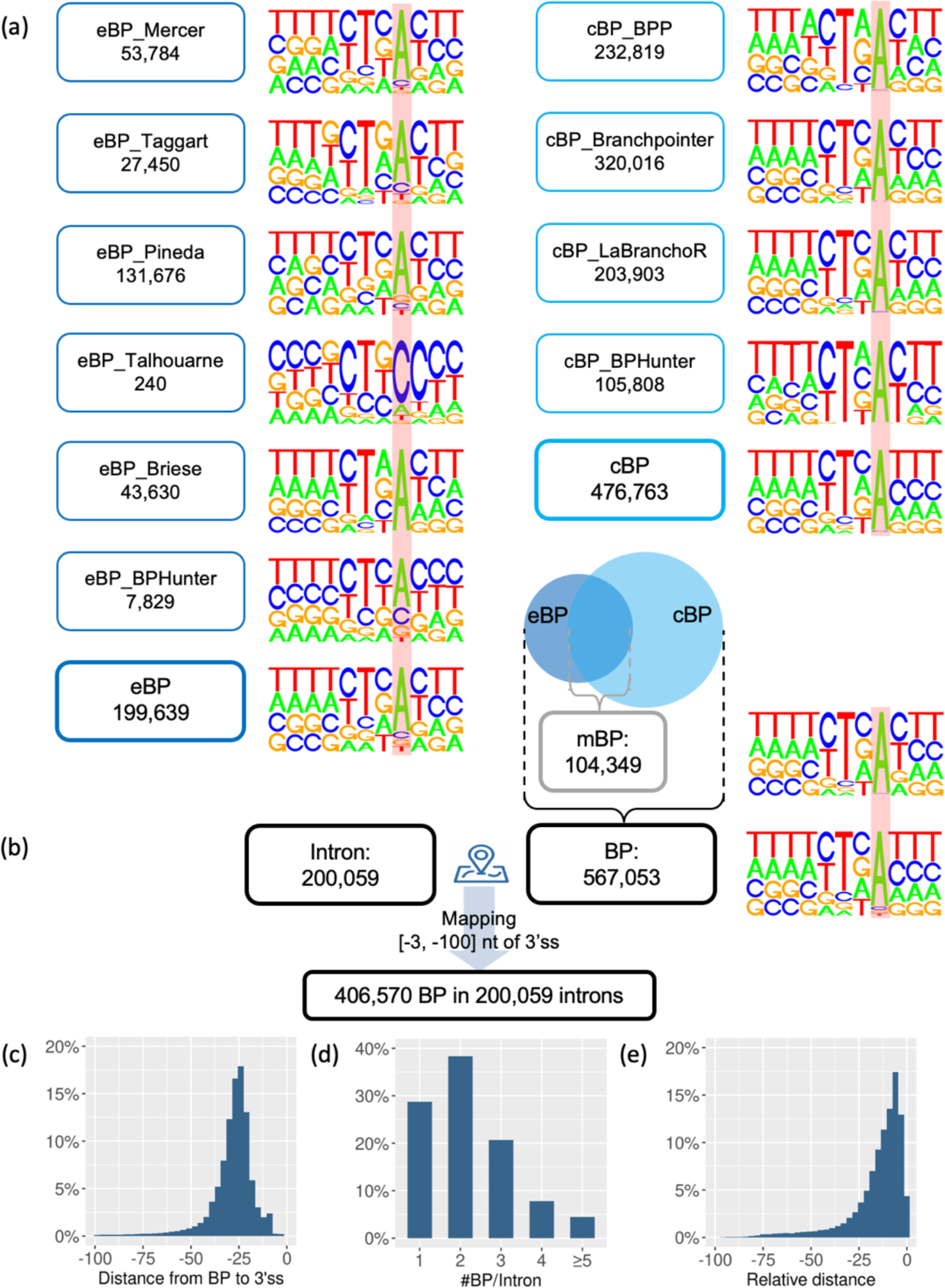
Integration of positional adjusted eBP and cBP, and mapping of BP onto introns. (a) The nucleotide composition displays the motif [-7, +3] of BP, where the locations of BP are marked with a red background. (b) BP were mapped to introns, focusing on the BP that resided within [-3, -100] nt upstream of 3’ss. (c) Distance from the mapped BP to their corresponding 3’ss. (d) The number of BP mapped to each intron. (e) The relative distance from the non-first BP to the first BP in each intron.

### Mapping BP to introns

We identified 200,059 unique introns from 41,975 transcripts of 17,372 protein-coding genes (Frankish et al. 2019), and classified introns into 199,393 (99.7%) major and 666 (0.3%) minor types (**Figure S3, Supplemental Data 2**). Because the major and minor spliceosomes regulate splicing differently (Turunen et al. 2013) (**Figure 1b**), it is important to distinguish intron types when studying BP and splicing. Major introns were dominated by GT-AG (98.9%) as the 5’ss-3’ss terminal dinucleotides, whereas 71% and 24% of minor introns were GT-AG and AT-AC, respectively. We therefore mapped 567,053 BP onto 200,059 introns, retained the BP within [-3, -100] nt of 3’ss, and finally obtained 406,570 BP associated with 200,059 introns (**Figure 3b, Supplemental Data 3** and **4**). We excluded the distal BP in deep introns in this study, as more complicated splicing mechanisms may occur (e.g., recursive splicing for multi-step intron removal (Sibley et al. 2015; Wan et al. 2021), stem-loop RNA structures to bring distal BP closer to 3’ss to facilitate splicing (Lin et al. 2016) (**Figure S1**)). Among the intron-associated BP, 53% of them were located within a 10-nt window [-21, -30] nt of 3’ss, and 84% were located within a 25-nt window [-15, -40] nt of 3’ss (**Figure 3c**). 29%, 38% and 21% of introns had one, two and three BP mapped respectively, whilst 4% of introns harbored more than four BP (**Figure 3d**). With 71% of introns containing multiple BP, we found that these multiple BP were largely located in close proximity. By taking the first BP in each intron (i.e., the one closest to the 3’ss) as the reference, 23% and 48% of non-first BP were found to be located within 5-nt and 10-nt upstream of the first BP (**Figure 3e**). In introns with multiple associated BP, the first BP could be the most important, as a previous study of sequence determinants of splicing showed that BP closer to 3’ss were much more likely to influence splicing efficiency (Rosenberg et al. 2015). The BP usage is also expected to be context-specific in alternatively spliced transcripts, but we do not have the requisite data to differentiate the usage. Thus, in this study, we considered the BP-intron associations in a general setting.

### Nucleotide composition, distance from BP to 3’ss, and binding energy between BP and U2/U12-snRNA

In major introns, BP motifs exhibited high conservation of A (93%) at pos:0 and T (74%) at pos:- 2, and high pyrimidine content (C/T) at pos:-2 (85%), pos:-3 (73%) and pos:+1 (68%). In minor introns, BP showed high conservation of A (95%) at pos:0 and T (80%) at pos:-2, and high pyrimidine content at pos:[-2, -5] and pos:+1 with frequencies 90%, 78%, 68%, 65% and 69% respectively (**Figure 4a**, **Table S4**). We measured the distance from BP to 3’ss, and found 14% and 53% of BP in major introns were distributed in regions [-11, -20] and [-21, -30] nt of 3’ss, whereas 35% and 32% of BP in minor introns were found within these two regions (**Figure 4b, Table S5**). BP in minor introns were generally located closer to 3’ss than BP in major introns, which is consistent with the major spliceosome requiring the binding of *trans*-element U2AF65 between BP and 3’ss that needs a larger spacing, which is absent in the minor spliceosome (Turunen et al. 2013). We further estimated the binding energy between BP and snRNA (**Methods**), and found that BP in major introns displayed weaker binding with U2-snRNA than that between BP and U12-snRNA in minor introns (**Figure 4c, Table S5**). This may imply that the BP-U2 interaction in the major spliceosome can be complemented by other spliceosomal elements (e.g., SF3B1, SF3B6, PHF5A, SFA2, SFA3) (Tholen et al. 2022), hence a weaker wobble BP-U2 base-pairing could be acceptable. Since the minor spliceosome forms a U11-U12 di-snRNA complex to bind to 5’ss and BP simultaneously, it requires more perfect BP-U12 base-pairing, which is also reflected in more conserved 5’ss and BP motifs in minor introns (**Figure 4a**). We further examined the region [-50, +20] nt of BP for motif enrichment (Bailey et al. 2009), and reported 17 enriched motifs (*p*-values < 0.05 and > 5% occurrence) in the 50-nt upstream region of BP in major introns (**Figure S4**). These motifs could represent potential *cis*-acting elements with a role in facilitating splicing. The co-occurrence between some motifs and BP might suggest different modes of collaboration between splicing factors.

**Figure 4:**
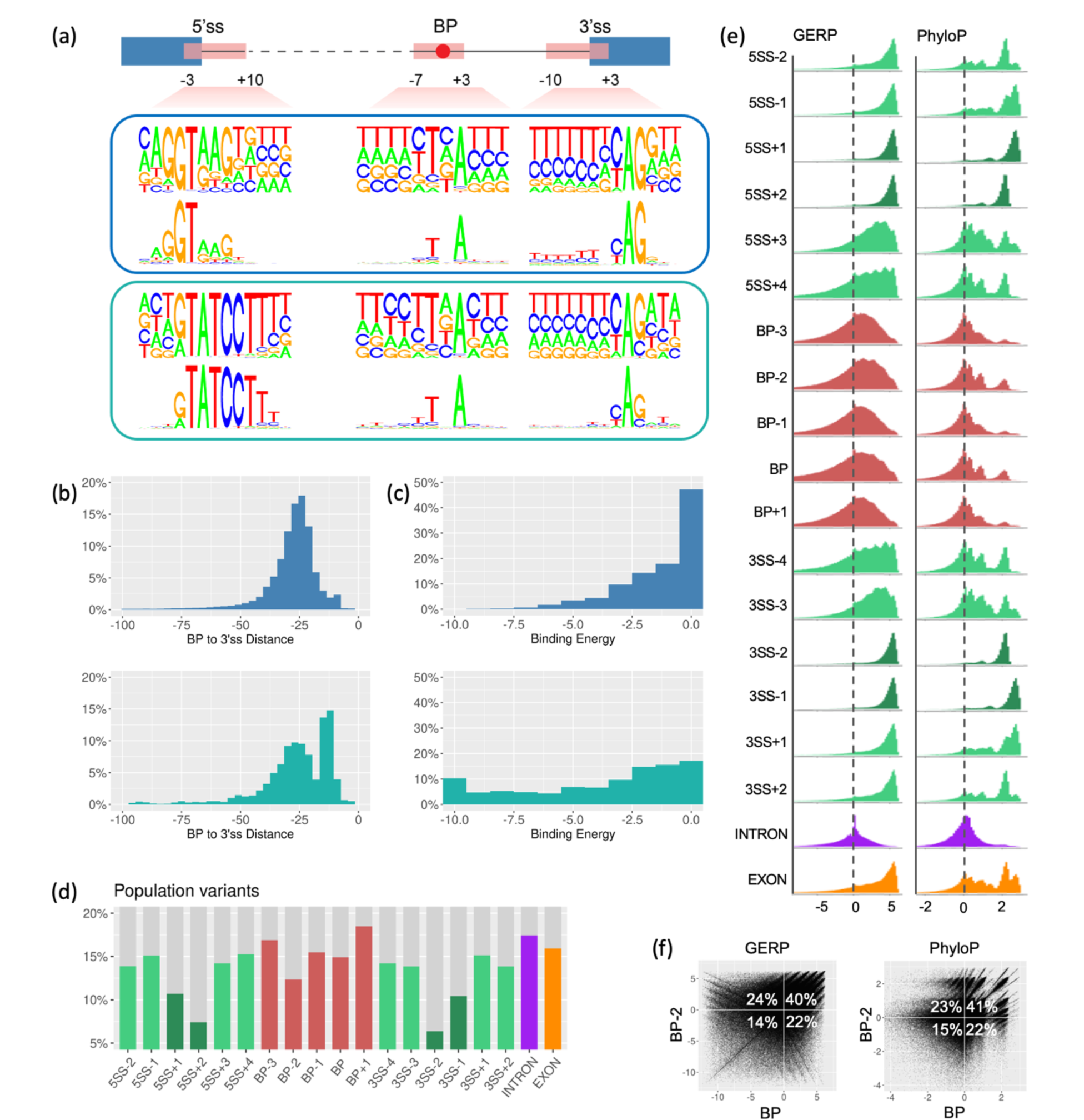
Characterization of BP. (a) Nucleotide composition of BP motifs, 5’ss motifs and 3’ss motifs in major and minor introns respectively, each box displaying the nucleotide frequency (upper) and information content (lower) of these motifs. (b) The distance from BP to 3’ss in major and minor introns respectively. (c) The binding energy between BP motif and U2/U12 snRNA in the major and minor introns respectively: the lower the energy, the higher the binding affinity. (d) The proportion of each genomic position harboring human population variants. (e) The distribution of the conservation scores GERP (left) and PhyloP (right) in each genomic position. (f) The cross-compared conservation scores between BP and BP-2 positions.

### Population variants in BP

We obtained human population variant data from the gnomAD database (Karczewski et al. 2020) r3.1.1, with which to compare the occurrence of population variants in the BP region, essential/flanking splice sites, and intronic/exonic background. We focused on the [-3, +1] positions of 406,570 BP, took a 6- nt window for each splice site ([-2, +4] of 5’ss and [-4, +2] of 3’ss) from 200,059 introns, and randomly sampled 1,000,000 positions from the intron and exon backgrounds. In this way, we generated 19 sets of genomic locations for comparison of their population variants (**Figure 4d, Table S6**). We found that 17.4% of intronic background and 15.9% of exonic background positions harbored population variants; the flanking splice sites had 13.9% - 15.3% of population variants, whereas the essential splice sites exhibited a much lower proportion (6.4% - 10.7%) of population variants. With regard to the BP region, 14.9% of BP positions, 15.5% of BP-1 positions, and 12.4% of BP-2 positions harbored population variants, which were lower than those for the intron and exon backgrounds, and similar to the flanking positions of splice sites. The other two positions in the BP region (BP-3 and BP+1) had 16.9% and 18.5% of population variants, which were close to or even higher than the random intronic positions. These findings suggested that the BP and BP-2 positions exhibit a lower rate of variation than the intronic background in human populations and are similar to the exonic background and flanking positions of essential splice sites.

### Cross-species conservation of BP

We obtained genome-wide conservation scores, GERP (based on 35 mammals (Cooper et al. 2005)) and PhyloP (based on 46 vertebrates (Pollard et al. 2010)) (**Methods**), and evaluated the level of evolutionary conservation in the abovementioned 19 sets of genomic locations. A positive score denotes that the position is likely to be evolutionarily conserved whereas a negative score indicates that the position is probably evolving neutrally (**Figure 4e, Table S7**). Both conservation scores were found to be comparable for each position studied. The essential splice sites and their exonic flanking regions displayed an absolute positive distribution of conservation scores, whereas their intronic flanking sites exhibited a reduced level of conservation. Intronic positions were mostly around zero whereas exonic positions were mostly positive. The BP region displayed a larger proportion of positive scores (evolutionarily conserved) than negative scores (evolving neutrally), with two peaks of high PhyloP scores for BP and BP-2. We cross-compared the scores between BP and BP-2 (**Figure 4f**), and found that ∼24% of negatively scored BP positions had positive scores for their corresponding BP-2 positions (2^nd^ quadrant), whilst ∼22% of negatively scored BP-2 positions had positive scores for their corresponding BP positions (4^th^ quadrant). Thus, if we accept at least one positive score for BP or BP-2 position, 86% of BP regions were evolutionarily conserved. Taken together, these findings suggested that the BP region is more conserved than the intronic background, but less conserved than the exonic background and splice sites; they also hint at a possible collaborative role between BP and BP-2 to ensure a certain level of conservation over evolutionary time. A recent study of the bovine genome showed that bovine BP are under evolutionary constraint (Kadri et al. 2021). It is worth noting that splicing evolves rapidly between species (Keren et al. 2010), with one study showing significant splicing differences between three primates, including human and chimpanzee (Xiong et al. 2018). This may suggest that the conservation scores of splicing regulatory elements may not invariably be correlated with their functional importance. In other words, BP with low/negative conservation scores could be functionally important.

### Comparative characterization of BP

We made seven comparisons of BP in the major introns by different definitions, from intron-level, to transcript-level, gene-level and chromosome-level. (I) 375,002 adenine-BP *vs*. 13,827 cytosine-BP *vs*. 9,349 guanine-BP *vs*. 6,635 thymine-BP (**Figure S5**). We found that adenine-BP and thymine-BP had much lower rates of population variation than cytosine-BP and guanine-BP (which reached ∼24%, considerably higher than the intronic background of 17.4%). Two spikes of high PhyloP conservation scores were observed for adenine-BP and thymine-BP. (II) 95,814 mBP *vs.* 56,547 exclusively eBP *vs.* 252,452 exclusively cBP (**Figure S6**). mBP displayed the most conserved nucleotide composition, the lowest rate of population variation, and the largest proportion of positive conservation scores, followed by exclusively cBP and then exclusively eBP. (III) 199,393 first BP in introns *vs.* 205,420 non-first BP in introns (**Figure S7**). Apart from the distance to 3’ss (median: -22 *vs.* -31 in first *vs.* non-first BP), which was determined by the definition of their group, no distinct feature was noted in this comparison. (IV) 57,418 BP in introns with single BP *vs.* 347,395 BP in introns with multiple BP (**Figure S8**). BP in introns containing a single BP showed an absolute dominance (96%) of T in the BP-2 position, a lower rate of population variation, and a larger proportion of positive conservation scores, than BP in introns with multiple BP. (V) 353,019 BP in introns without alternative splicing *vs.* 51,794 BP in introns with alternative splicing (**Figure S9**). No notable difference was observed. (VI) 156,413BP in genes encoding a single isoform *vs.* 248,400 BP in genes encoding multiple isoforms (**Figure S10**). No notable difference was observed. (VII) 14,834 BP in genes located on the sex (X and Y) chromosomes *vs.* 389,979 BP in genes located on the autosomes (**Figure S11**). The sex chromosomes had a slightly higher proportion of introns with single BP (32% *vs.* 28%) than the autosomes. BP on the sex chromosome also exhibited slightly higher conservation scores. We observed only 8.7% of BP on sex chromosomes harboring population variants. By contrast, 15.1% of BP on the autosomes harbored population variants, revealing a much lower rate of population variation of BP on the sex chromosomes than the autosomes.

### Pathogenic BP mutations

Mutations in BP may lead to reduced, altered or abolished binding to spliceosome elements during the beginning of splicing, leading to the disruption of splicing and giving rise to mis-spliced gene products with the consequent loss of expression/function (**Figure 1c**). Through HGMD and the primary literature, we identified a total of 56 pathogenic BP mutations (48 single-nucleotide variants (SNVs) and 8 micro- deletions) in 44 human genes underlying 40 different inherited disorders (**Table 2**), all of which had been experimentally confirmed by analysis of the mis-splicing consequences. These disorders were attributed to different classes of disease: developmental, oncogenic, metabolic, neurological, immunological and circulatory. These BP mutations (47 germline and 9 somatic) comprised 16 homozygotes, 22 heterozygotes, 6 hemizygotes and 12 without zygosity information, and were located within [-9, -40] nt of 3’ss. Taking the reported mis-splicing consequences together (and bearing in mind that any one mutation may have multiple mis-splicing consequences at the RNA level), we identified 37 complete and 7 partial exon skippings, and 10 complete and 11 partial intron retentions, suggesting that complete exon skipping may be the predominant molecular consequence of the disruption of BP recognition. The discovery of these pathogenic BP mutations generally occurred in a similar context (**Box 1**), while two recent studies used some of the published BP datasets to study cancer genes (Leman et al. 2020; Canson et al. 2021). Their discovery timeline showed that 11, 15 and 30 pathogenic BP mutations were published in the 1990s, the 2000s and since 2010, respectively. Thus, neither the advent of NGS technology (since 2010), nor the publication of large-scale human BP datasets (since 2015), has resulted in a major increase in the identification of pathogenic BP mutations. In fact, the proportion of pathogenic BP mutations among all reported pathogenic mutations has been consistently very low since the 1990s. Based on HGMD, the proportion of BP mutations among all reported lesions is currently 56/323,661=0.00017; however, it is hard to escape the conclusion that the discovery of BP mutations has hitherto been largely serendipitous. It follows that the detection efficiency of pathogenic BP mutations is probably rather low, when compared with other types of lesions. Therefore, there is an urgent need to bridge BP data with NGS data, thereby extending the utility of the NGS data, making more practical use of the BP data, and contributing to pathogenetic discovery.

**Box 1:**
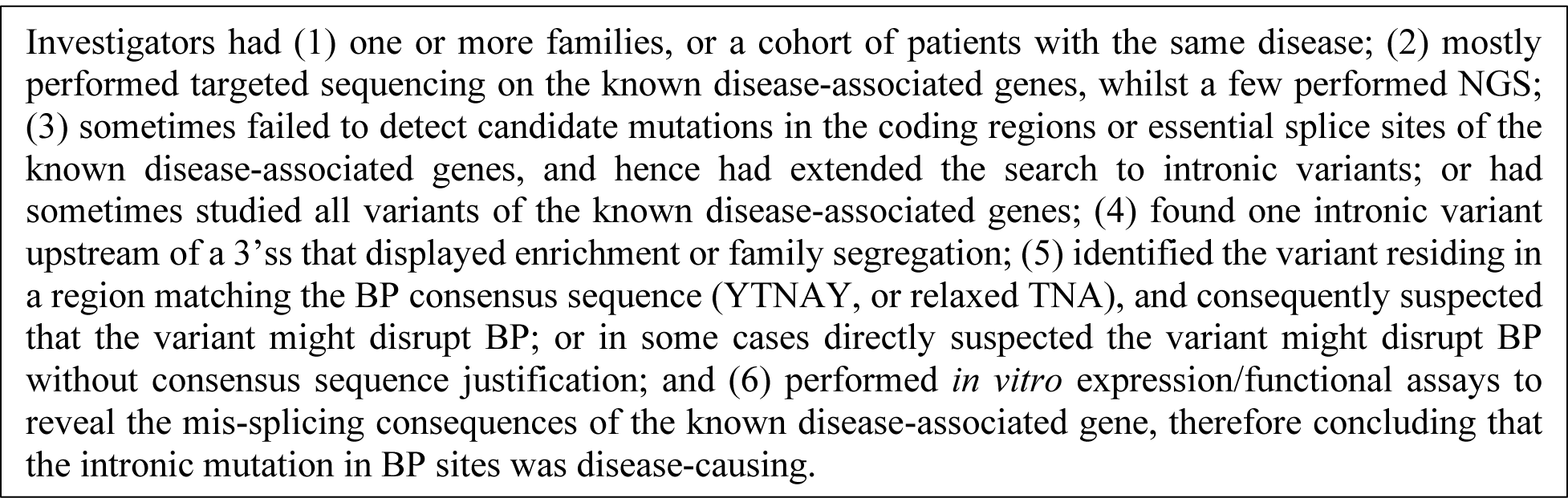
The typical discovery narrative of the published pathogenic BP mutations.

**Table 2:**
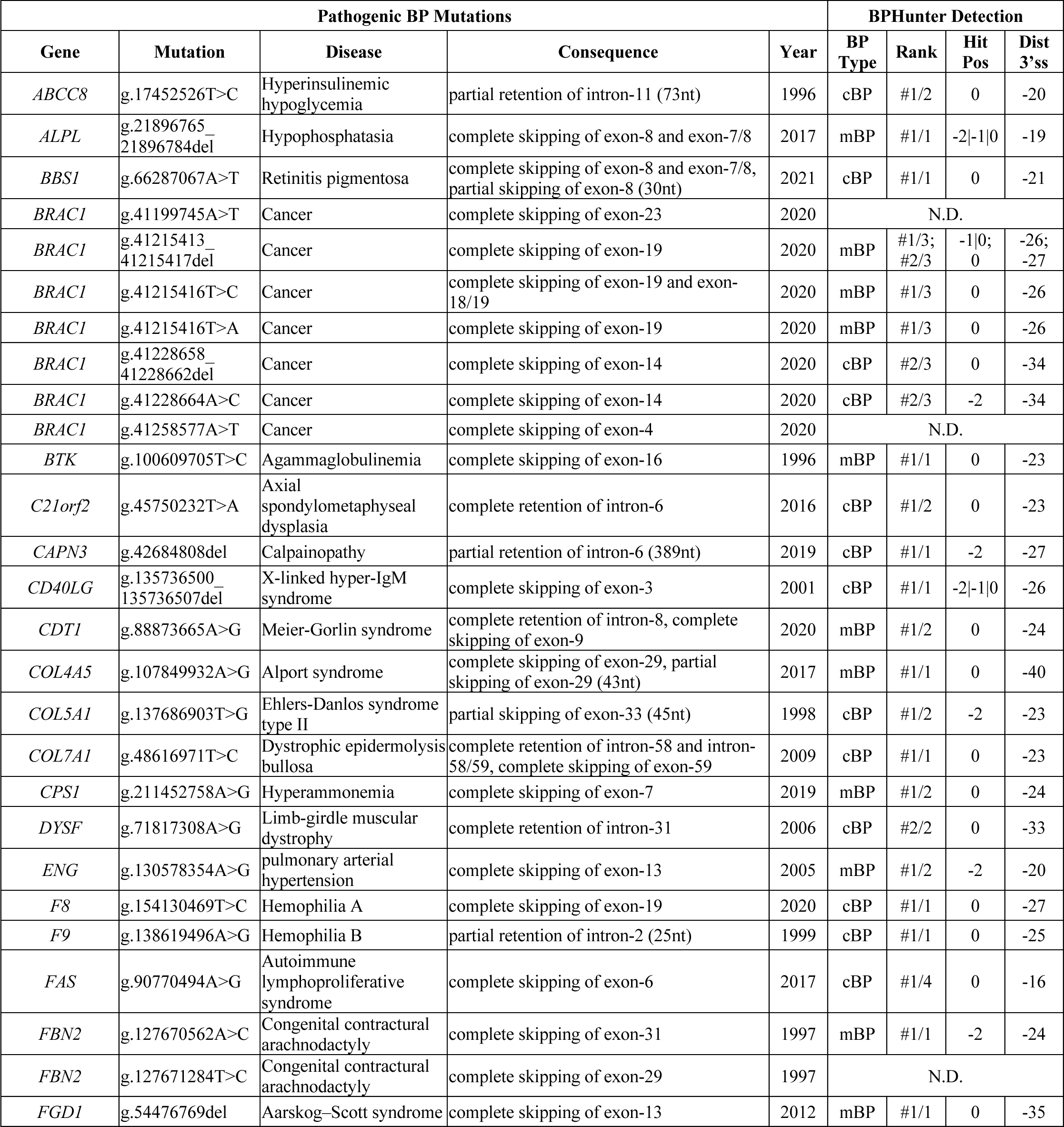
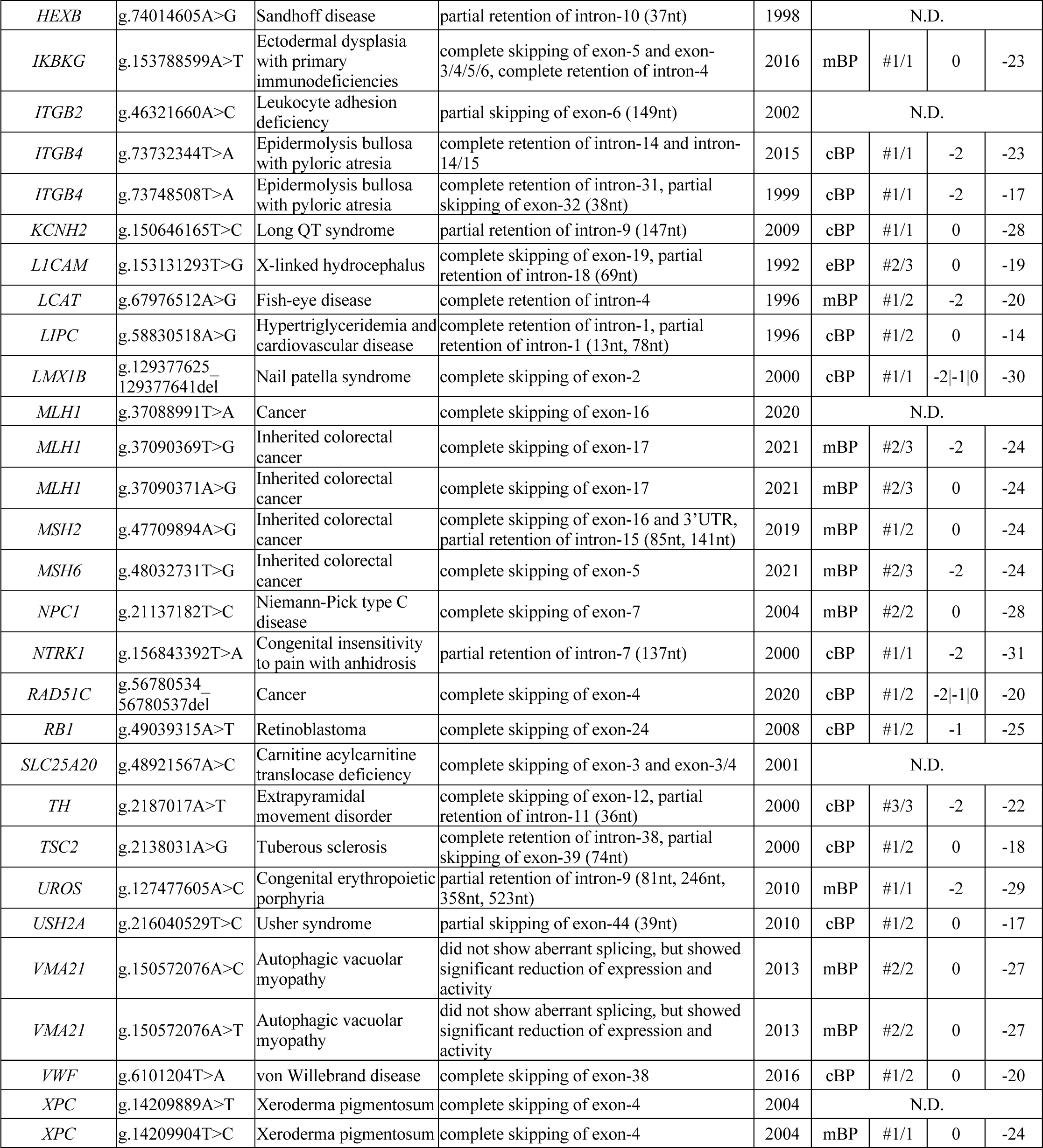
The 56 reported pathogenic BP mutations in 44 genes underlying human inherited disorders, with experimentally confirmed molecular consequences. BPHunter successfully detected 48 of them. Full information and references are available in Supplemental Data 5. (The genomic positions are on hg19/GRCh37 human genome assembly. Zygo: zygosity. Hit Pos: BP positions disrupted by the variant. N.D.: not detected.)

### BPHunter: detection of intronic variants that disrupt the branchpoint

Variants that affect BP recognition may be individually rare, but their impact on splicing, gene expression, and protein function can be highly deleterious. Therefore, in searching for disease-causing variants, we should routinely consider BP mutations as a discrete category of potentially damaging variants, especially when the search within coding regions and essential splice sites has turned out to be unproductive. We therefore developed BPHunter for the genome-wide detection of human intronic variants that disrupt the branchpoint in NGS data, thereby enabling the identification of BP mutation candidates that could potentially result in aberrant splicing at a biochemical level and hence disease at a physiological level. Given a VCF file of variants, BPHunter detects those variants that disrupt BP with informative outputs, including gene, transcript, intron, BP name, BP ranking, BP positions disrupted, distance from BP to 3’ss, population variant MAF, conservation scores, supporting sources, etc. (**Data and Web Access**). We developed the BPHunter program as a one-line command that can be easily implemented into NGS analysis, and also as a user-friendly webserver for users with less computational expertise. We believe that there may have been many negative findings involving BP variants that were tested but never reported. To improve our understanding of functional BP, their mutational consequences, and their cell-type specificity, we ought to extract value from these negative data. Therefore, we created a reporting system in the BPHunter webserver for researchers to submit their tested BP variants that do not display aberrant splicing. These data will be periodically collated, analyzed and deposited as a preprint, with all contributing researchers acknowledged as members of the BPHunter Effort Group. As we cannot validate the submitted negative BP variants centrally, we shall require all investigators to take responsibility for their submitted data. In summary, BPHunter constitutes a computational method capable of detecting candidate BP-disrupting mutations, and a platform for collating the negative results from BP genetics studies that should not be wasted.

### Validation of BPHunter on the reported pathogenic BP mutations

We deployed BPHunter retrospectively on the 56 reported pathogenic BP mutations, and successfully captured 48 of them (**Figure 5a, 5b** and **S12, Table 2, Supplemental Data 5**). Six deletions (4/5/5/8/17/20-nt) removed the entire/partial BP motif, two 1-nt deletions removed one BP site and one BP- 2 site respectively, 27 SNVs disrupted BP sites, 12 SNVs disrupted BP-2 sites, and one SNV disrupted a BP-1 site (**Figure 5c**). In 19 cases, the mutations disrupted the only BP in their introns; in 18 cases, the mutations disrupted the first BP in introns with multiple BP (14 introns with two BP, three introns with three BP and one intron with four BP); and in 11 cases, the mutations disrupted the non-first (10 second and one third) BP (**Figure 5d**). The affected BP comprised 22 mBP, one exclusively eBP and 25 exclusively cBP (**Figure 5e**). There were 25 BP that matched the YTNAY consensus sequence, whereas the other 8, 12 and 3 BP matched the more relaxed YTNA, TNA and YNA consensus motifs, respectively (**Figure 5f**). There were 41 BP positions supported by more than one source (**Figure 5g**). No population variants were found for 43 BP positions, whereas very rare variants were noted in the other five BP positions (**Figure 5h**). The conservation scores GERP and PhyloP were both positive for 39 BP positions and both negative for seven BP positions (**Figure 5i**), whilst 9 of the 11 mutations disrupting the non-first BP had both positive scores. In addition, we deployed two popular mis-splicing prediction tools (SpliceAI (Jaganathan et al. 2019) and MMSplice (Cheng et al. 2019)) and a mutation deleteriousness prediction score (CADD (Rentzsch et al. 2019)) to evaluate these pathogenic BP mutations but none of them performed well (**Figure 5j**). SpliceAI, providing a probability score for altering the acceptor sites, predicted only one mutation above its high-precision cutoff of 0.8, and eight mutations above its recommended cutoff of 0.5. MMSplice predicted only one mutation beyond its suggested acceptor-disruption cutoff of -2. CADD, a composite mutation damage score which is heavily reliant on conservation-related metrics, identified 3 mutations having scores >20 (top 1%), 35 mutations having scores in the range 10-20 (top 1-10%), and 10 mutations with scores <10.

**Figure 5:**
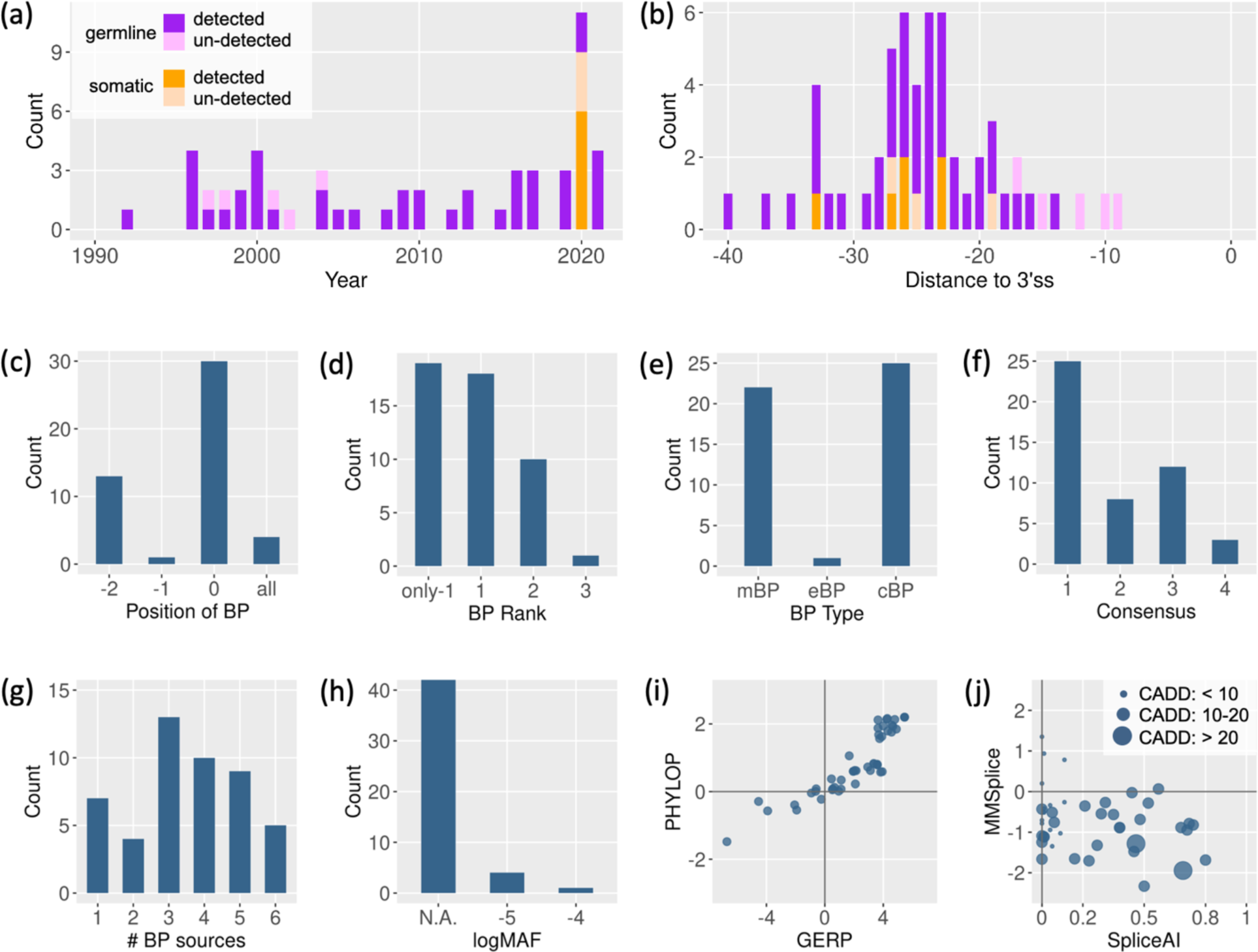
Detection of pathogenic BP mutations by BPHunter. (a) Timeline of the 56 published reports of pathogenic BP mutations. (b) Distance from the reported BP mutations to their 3’ss. (c) Disrupted positions of BP. (d) Ranking of the disrupted BP in its intron. (e) Type of disrupted BP. (f) Matched consensus sequence of the disrupted BP (1: YTNAY, 2: YTNA, 3: TNA, and 4: YNA). (g) Number of data sources supporting the disrupted BP. (h) Population variation MAF of the disrupted BP. (i) Conservation scores (PhyloP *vs.* GERP) of the disrupted BP. (j) Mis-splicing scores (MMSplice *vs*. SpliceAI) and the mutation deleteriousness score (CADD) of BP mutations.

### Pathogenic BP mutations that BPHunter did not detect

BPHunter retrospectively detected 48/56 reported pathogenic BP mutations, leaving eight BP mutations undetected (**Table 2**). Five were published around the year 2000 and were located very close (- 9/-10/-12/-15/-17 nt) to 3’ss, whereas three were published in 2020 and were located -19/-25/-27 nt to 3’ss (**Figure 5a** and **5b**). The eight introns harboring these mutations all had documented BP sites in our database (three introns with single BP, and five introns with two BP), but none of these pathogenic mutations disrupted the BP/BP-2 sites in our database (**Figure S13**). By extracting and aligning the wt and mt sequences from these eight mutations using SeqTailor (Zhang et al. 2019), we found that all of them created an AG dinucleotide in the region between BP and 3’ss (**Figure S13**). By comparing the MaxEnt 3’ss strength (Yeo and Burge 2004) between the constitutional AGs and the newly created AGs, we found that these eight mutations created one stronger 3’ss, five weaker 3’ss, and two unlikely 3’ss, respectively. We therefore reasonably hypothesize that the AG dinucleotides created in the region between BP and 3’ss may promote binding with the *trans*-element U2AF35 that recognizes the acceptor site, thereby shortening the original spacing from BP to the constitutional AG that would normally be bound by another *trans*-element, U2AF65 (**Figure 1b**). In these cases, the structural organization and spacing requirement of the intron were disrupted, thereby disrupting the constitutional splicing events. Thus, these eight mutations should not be considered as BP mutations. They presumably cause aberrant splicing through the disruption of another important process in the splicing machinery. BPHunter therefore successfully detected 100% of the reported pathogenic BP mutations, in an efficient, systematic and informative manner.

### Comparison of the pathogenic BP mutations with common BP variants in the general population

Focusing on SNV, we compared the BP sites harboring 38 pathogenic mutations (26 mutations disrupting BP and 12 mutations disrupting BP-2) against BP sites harboring 1,288 common variants in population (730 variants on BP and 558 variants on BP-2, with MAF > 5%) (**Figure S14a**). We found that common variants occurred significantly less frequently in introns with single BP (*p*-value < 0.0001, versus pathogenic mutations, by Fisher’s Exact test), and were significantly more frequent in BP of rank≥#3 (*p*- value < 0.0001). Some 34% of common variants were located within BP motifs that did not match any level of consensus sequence (*p*-value < 0.0001), whilst 61% of common variants were located in BP sites supported by only one source (*p*-value < 0.0001). The variants in BP and BP-2 positions were then analyzed separately. We compared 26 pathogenic BP mutations against 730 common BP variants (**Figure S14b**). The nucleotide change A>G was the most frequently encountered (73%) among the pathogenic BP mutations, and was marginally significant when compared with common BP variants (*p*-value = 0.028). We also compared 12 pathogenic BP-2 mutations against 558 common BP-2 variants (**Figure S14c**). The nucleotide changes T>A and T>G in pathogenic BP-2 mutations were both significantly enriched against common BP-2 variants (*p*-value = 0.0358 for T>A, and *p*-value = 0.0005 for T>G). The nucleotide change T>C was the least frequent (17%) among pathogenic BP-2 mutations although the most frequent (36%) among common BP-2 variants, albeit not significant (*p*-value = 0.1471). The conservation scores for both BP and BP-2 positions did not exhibit any differences between pathogenic mutations and common variants. However, statistical validation was based on a relatively small number of reported pathogenic mutations shown to disrupt BP, and once additional BP mutations have been reported, this statistical analysis should be refreshed.

### Strategy for prioritizing BP mutation candidates

The in-depth analysis of the published pathogenic BP mutations, in concert with the analysis of common variants in the BP region, helped us to formulate a strategy to prioritize BP mutation candidates: (1) deletion of the entire BP motif, or disruption of the BP or BP-2 positions; (2) mutation of the only BP in an intron, or the first/second BP in an intron with no more than three BP; (3) mutation of BP motifs matching the consensus YTNAY, whilst also allowing slightly relaxed consensus; (4) mutation at BP positions with more than one supporting source, irrespective of the type of source (mBP/eBP/cBP); (5) mutation at BP positions with no or rare population variants; and (6) mutation at BP positions with positive- scored GERP or PhyloP may be preferred but not invariably. The computation of SpliceAI, MMSplice and CADD scores could be helpful to provide a reference, but should not be relied upon, particularly when evaluating newly identified BP mutations (see our case study of a *STAT2* mutation in a critical COVID-19 patient). A high prediction score cutoff could lead to the loss of true positives and a lowered prediction score cutoff could lead to a large number of false positives. Additionally, homozygous and hemizygous variants should be prioritized, candidate mutations in genes associated with the disease under study should be prioritized, and the NGS sequencing quality needs to be taken into consideration. This prioritization strategy will be updated as the number of discovered BP mutations increases.

### A germline heterozygous single-nucleotide variant of *STAT2* disrupts BP and splicing in a patient with critical COVID-19

We performed BPHunter prospectively on whole-exome sequencing (WES) data from a cohort of 1,035 patients with critical COVID-19 (Casanova et al. 2020), focusing on 13 genes that have been reported to carry deleterious variants impairing type-I IFN production (Zhang et al. 2020). BPHunter detected one private heterozygous variant g.56749159T>A (IVS5-24A>T) in *STAT2* that disrupted the only BP (mBP) located -24 nt from 3’ss of intron-5 (**Figure 6a**). This BP matched the YTNAY consensus sequence, had a MAF of 6.98e-6, exhibited a GERP of -0.257 and PhyloP of 0.152, and was supported by five sources. These features satisfied our prioritization strategy requirements for a promising candidate BP mutation even though the mis-splicing/deleteriousness predictions were not suggestive of its pathogenicity (SpliceAI = 0, MMSplice = -0.4, CADD = 10.1). We therefore hypothesized that this intronic variant might disrupt BP recognition leading to aberrant splicing of *STAT2* transcripts. To assess its impact on mRNA splicing, we performed an exon trapping assay. Compared to the wild-type control, mRNA extracted from cells transfected with this variant displayed a majority of abnormally spliced *STAT2* transcripts: 6% normal transcripts in the mutant *vs.* 70% in the control, 82% of complete exon-6 skipping in the mutant *vs.* 25% in the control, and 4% of complete intron-5 retention in the mutant *vs.* none in the control (**Figure 6b**). We extracted mRNA from whole blood of the patient and a healthy control, and amplified the *STAT2* cDNA from exon-3 to the exon-7/8 boundary. Sanger sequencing of the PCR products following TOPO-TA cloning then showed that the patient’s cells had more mis-spliced transcripts (exon skipping and intron retention) than the healthy control (**Figure 6c**). The cDNA from the patient contained 35% transcripts that were not annotated in GTEx, whilst the control contained 9% of unannotated transcripts (**Figure 6d**). We also assessed *STAT2* mRNA by RT-qPCR to estimate the amount of canonical mRNA transcripts extracted from whole blood, and observed about half the level in the patient as compared with the control (**Figure 6e**). If we also consider the mis-spliced gene products that could have been degraded by NMD (Hauser et al. 2020), the patient’s cells should carry more aberrantly spliced products. This case study effectively demonstrated the identification of a novel BP mutation in *STAT2* from a life-threatening COVID-19 patient, and the experimental validation of its biochemical consequences. It also showed that a BP position characterized by a very low level of evolutionary conservation can nevertheless be functional in splicing regulation and it is for this reason that existing splicing/deleteriousness prediction tools are inadequate to the task of identifying candidate BP mutations.

**Figure 6:**
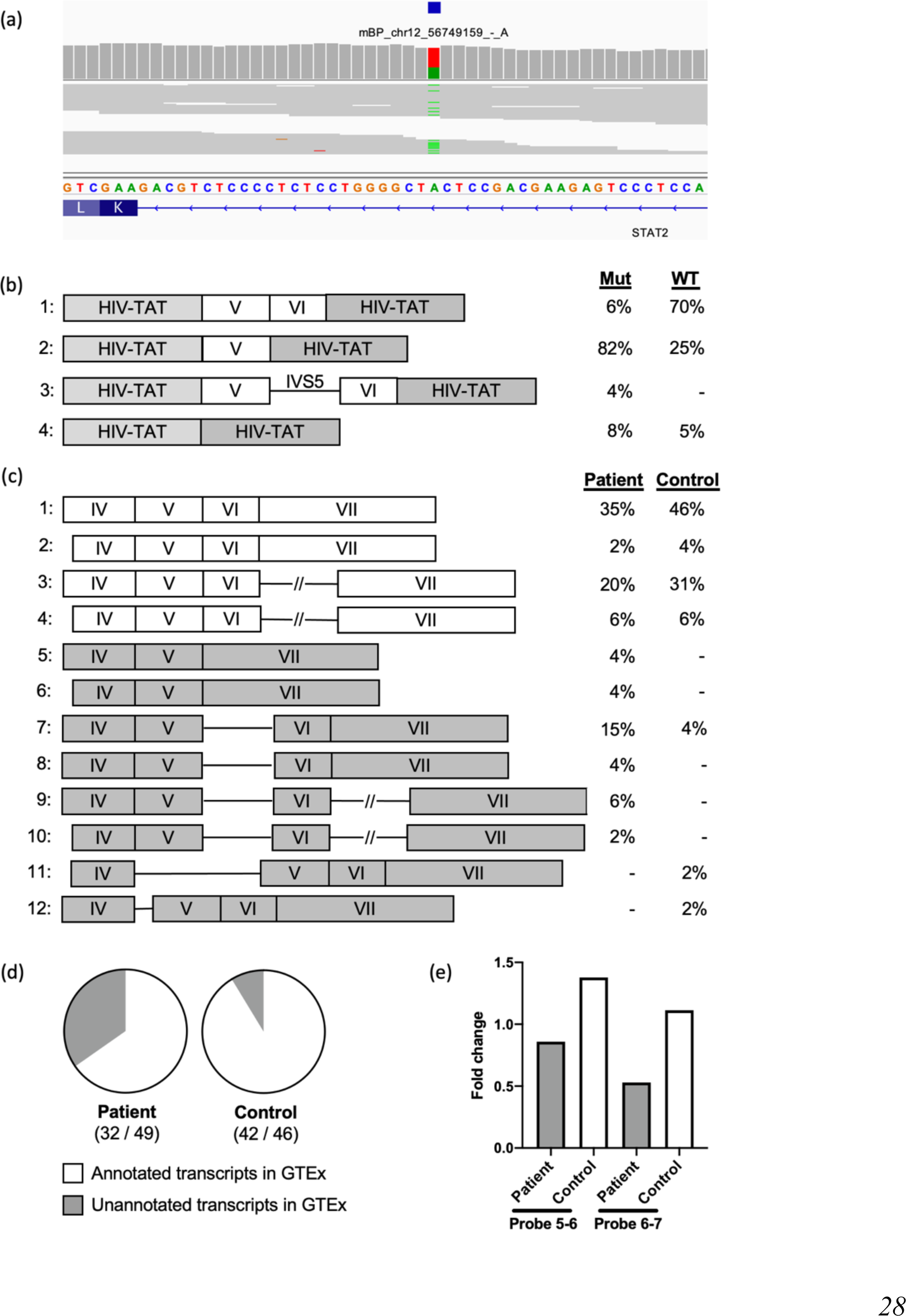
Detection and validation of a private heterozygous *STAT2* variant disrupting BP and splicing in a life-threatening patient with COVID-19. (a) Detection of a private heterozygous *STAT2* variant disrupting BP. (b) *STAT2* transcripts and proportions from an exon trapping assay in COS-7 cells. (c) *STAT2* transcripts and proportions from RNA extracted from whole blood. (d) Ratio of the annotated transcripts in GTEx versus the total number of transcripts. (e) Estimation of *STAT2* canonical transcripts based on RT-qPCR, measuring *STAT2* mRNA levels in whole blood using two probes spanning intron 5 (probe 5-6) and intron 6 (probe 6-7), respectively.

### A somatic intronic 59-nt deletion of *ITPKB* removes all three potential BP sites and leads to the retention of the intron in a lymphoma patient

We further performed BPHunter on somatic tumor-specific mutation data derived from the whole-genome sequencing (WGS) of 53 diffuse large B-cell lymphoma (DLBCL) tumor samples (Georgiou et al. 2016; Ren et al. 2018; Ye et al. 2021). Focusing on a set of 212 lymphoma-associated genes (Ren et al. 2018), BPHunter detected one intronic deletion in *ITPKB,* from one DLBCL sample, that removed 59 nucleotides from intron-3 (g.226835095_226835153del) (**Figure 7a**). This intron contains three potential BP sites, and the deletion eliminated all three of them, suggesting that the *ITPKB* gene product is highly likely to be mis-spliced in this tumor sample. *ITPKB* has been shown to be a negative regulator of the BCR/NF-κB signaling pathway which is very important for the growth and survival of lymphoma cells (Schmitz et al. 2018). It has also been shown that mutant (truncated) ITPKB proteins boost PI3K/Akt signaling which is a key growth-promoting pathway in lymphoma cells (Tiacci et al. 2018). We therefore hypothesized that this intronic deletion would lead to aberrant splicing of this intron, resulting in a truncated and/or decayed *ITPKB* gene product with loss-of-function/expression, leading to promotion of the growth and proliferation of lymphoma cells in the affected patient. We then studied RNA-seq data of this tumor sample derived from the patient, as well as another three tumor biopsies from random DLBCL patients who were found not to carry this mutation, and observed complete retention of intron-3 specifically in this patient (**Figure 7b** and **7c**). Hence, we concluded that this intronic deletion, which would have been filtered out if without BPHunter, resulted in aberrant splicing of *ITPKB*, which consequently contributed to the lymphomagenesis in this patient. The mis-splicing predictions failed to suggest the pathogenicity of this intronic deletion. This case study successfully demonstrated the use of BPHunter for the detection and deeper exploration of somatic mutations. In addition, using BPHunter, we screened the non-coding somatic mutations documented in COSMIC (Tate et al. 2019), focusing on 1,148 selected genes known to be associated with cancer formation and progression (**Methods**), and identified 180 candidate BP mutations (133 SNVs and 47 deletions, in 126 genes, from 171 patients involving 25 cancer types) that appear to have the potential to disrupt splicing as cancer driver or passenger mutations (**Supplemental Data 6**).

**Figure 7:**
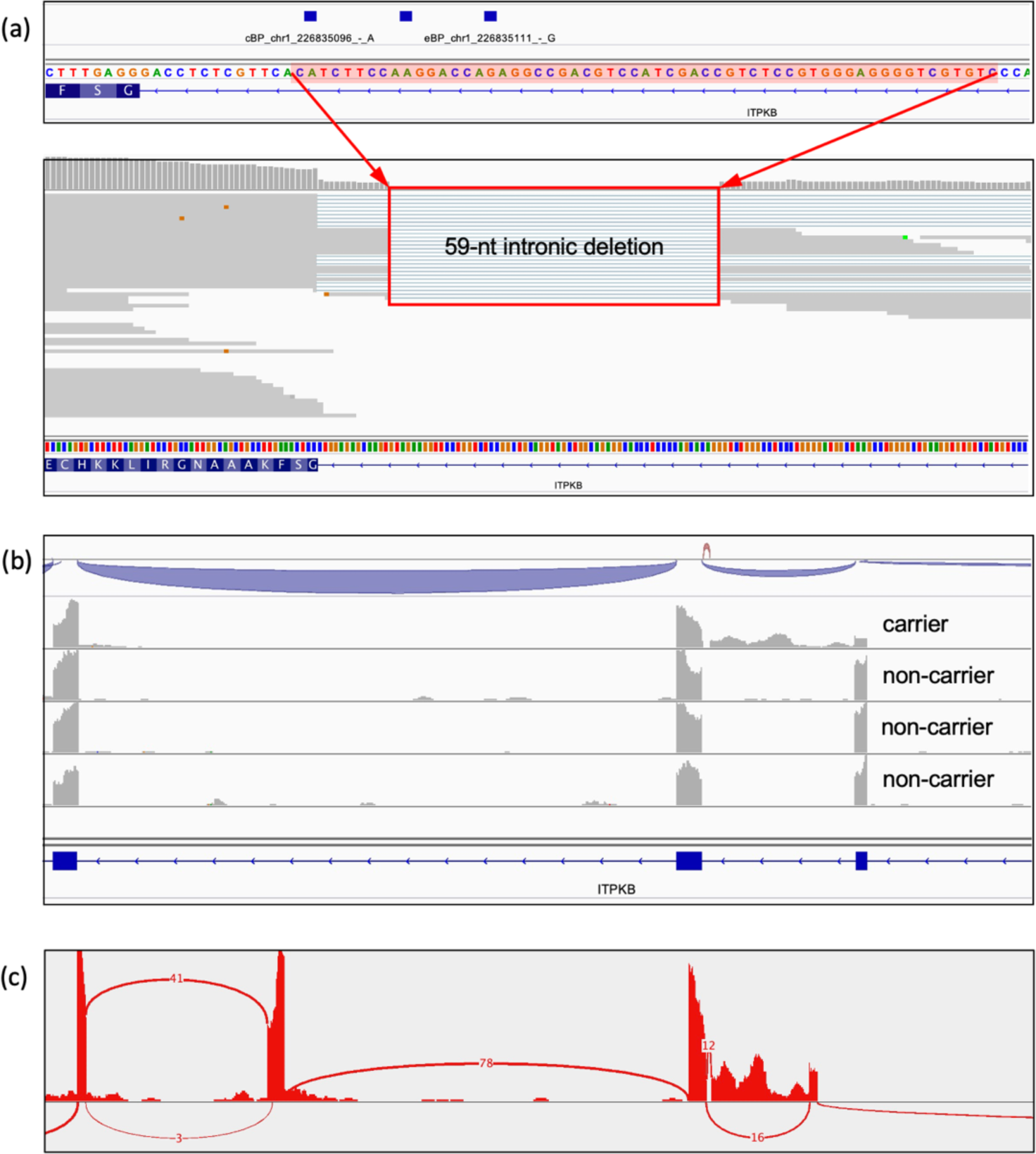
Detection and validation of a private somatic intronic deletion of *ITPKB* in a lymphoma patient. (a) Detection of a private somatic intronic 59-nt deletion of *ITPKB* that removed all three potential BP sites in intron-3. (b) RNA-seq reads alignment on *ITPKB* that showed intronic retention in the lymphoma patient carrying this mutation, but not in the patients without this mutation. (c) Sashimi plot from the RNA-seq data.

## DISCUSSION

Intronic sequences comprise ∼38% of the human genome, whilst exonic sequences comprise ∼3%. In the search for candidate mutations underlying human disease, current approaches mostly focus on non-synonymous variants in coding regions and variants residing in essential splice sites. If an investigator wishes to screen for candidate mutations among the vast number of intronic variants, a combination of approaches may be applied, e.g. testing for enrichment of intronic variants in a case-control study, computing and ranking the mutation deleteriousness/mis-splicing scores (Cheng et al. 2019; Jaganathan et al. 2019; Rentzsch et al. 2019) of intronic variants, referencing the MAF of population variations (Zhang et al. 2018a; Karczewski et al. 2020), and searching for genetic heterogeneity (presumably deleterious intronic variants together with coding/splice-site variants) underlying physiological homogeneity (Zhang et al. 2021). However, enrichment of intronic variants relies on having a group of carriers in cases versus controls, so individual variants (albeit functionally impactful) in sporadic cases are rarely captured. Moreover, many mutation deleteriousness scores (Rentzsch et al. 2019) are heavily reliant on evolutionary conservation, and splicing has been shown to evolve rapidly with significant differences between species (Keren et al. 2010; Xiong et al. 2018). This suggests that the application of conservation-derived deleteriousness scores to intronic splicing regulatory elements may not be particularly effective, a conclusion supported by our *STAT2* case study. We also showed that mis-splicing scores (Cheng et al. 2019; Jaganathan et al. 2019) are underpowered to prioritize promising BP mutations. Further, even though some intronic variant candidates may be identified, they do not come with the biochemical interpretations that other types of mutation do (e.g., missense, stop-gain, essential splice site), to guide the investigators in assessing their pathogenicity.

With BPHunter, we are now able to detect intronic variants that may disrupt BP recognition, and hence splicing and gene expression/function, in a systematic, efficient and informative manner. It provides an opportunity for disease genetics studies to explore patients’ NGS data more deeply, by identifying candidate BP mutations in known disease-associated genes, or alternatively in new and functionally promising candidate genes. It also provides an opportunity to systematically investigate the biomolecular effect of BP mutations, at least in some genes, particularly in those introns containing multiple BP. Such studies will delineate the intron-specific molecular associations between BP and splicing. We expect rapid expansion in the number of characterized BP mutations, and their biochemical consequences and pathophysiological impact, which will help us to comprehend BP function and dysfunction, from the positive results standpoint. BPHunter also provides a platform to report BP variants that do not give rise to mis-splicing, from the standpoint of negative results, which cannot be published but should nevertheless not be neglected. Lessons learnt from negative BP variants will feed back valuable input that can be used to determine functional BP in a cell-type context, which will in turn improve our understanding of BP and our ability to discover pathogenic BP mutations. Some studies proposed some BP variations without severe functional impact (Leman et al. 2020; Canson et al. 2021), and we expect to collect more such negative results through our platform for a larger-scale analysis.

We built a comprehensive database of human BP which represents a qualitative and quantitative knowledgebase of BP in the human genome. This BP database is however not yet optimal. The eBP are likely to include false-positive BP, caused by the mis-location of BP positions from RNA-seq reads. The cBP would have missed false-negative BP (some real non-adenine-BP for example), as machine learning methods always learn the major distinguishing features from training data to differentiate positives and negatives. Regarding the BP consensus sequence, we noted that while only ∼40% of BP motifs matched the long established YTNAY, most of the remaining BP motifs fitted with one of several increasingly relaxed consensus patterns. Our hypothesis-I is that BP motifs that fit the relaxed consensus (mostly displaying low binding affinity with snRNA) may have *cis*-elements (e.g., intronic splicing enhancers) in the vicinity that interact with *trans*-elements with stronger affinity, to guarantee the collaborative organization of the spliceosome necessary for correct splicing. Our hypothesis-II is that the definition of the BP consensus sequence could be context-specific (e.g., tissue-, cell-type-, developmental stage-, intron-specific, or RNA structure-dependent) to some extent, to meet the highly context-specific physiological requirements of different tissues and cells. In addition, we are aware of some splicing mechanism studies involving BP, including stochastic splice site selection that leads to kinetic variation in intron removal (Wan et al. 2021), recursive splicing that uses multiple intermediate 5’ss-BP-3’ss deep within the intron to allow multi-step intron removal (Sibley et al. 2015), stem-loop RNA structures that facilitate splicing by bringing distal BP closer to 3’ss (Lin et al. 2016), and the opening of the BP bulge which could occur during splicing (Konarska and Query 2005). These studies have presented even more complicated splicing mechanisms that call for greater effort in understanding the splicing process and in investigating the splicing-related pathogenesis.

BPHunter enables the efficient genome-wide detection of candidate BP mutations in NGS data, and we hope this tool will help to improve our ability to diagnose human genetic disease. We also anticipate the application of BPHunter to the study of human somatic mutations, in cancers for example (Group et al. 2020). Our timely recruitment of BPHunter to screen our COVID-19 cohort led directly to the identification of a novel and deleterious BP mutation in the *STAT2* gene from a life-threatening COVID-19 patient, which was immediately tested experimentally to demonstrate its biochemical consequences. Our subsequent application of BPHunter to lymphoma patients also led directly to the detection of an intronic deletion of *ITPKB* that removed all three potential BP sites in that intron leading to its complete retention as evidenced by RNA-seq data. Without BPHunter, neither of these deleterious mutations, that impaired the function of these key genes, would have been picked up by pre-existing classical methods (e.g., mis-splicing predictions, mutation deleteriousness scores, variant enrichment tests). BPHunter constitutes not only an important resource that provides us with a better genome-wide understanding of BP, but also a driving force that enables us to better discover BP variants underlying human disease.

## METHODS

### eBP data

We collected five datasets of experimentally identified high-confidence BP from five large-scale studies. We named these five datasets after the last names of their first authors: eBP_Mercer (Mercer et al. 2015), eBP_Taggart (Taggart et al. 2017), eBP_Pineda (Pineda and Bradley 2018), eBP_Talhouarne (Talhouarne and Gall 2018) and eBP_Briese (Briese et al. 2019) (**Table 1**). The first four datasets were derived from RNA-seq data: eBP_Mercer was identified from RNA-seq data from 11 cell lines (GSE53328) (Mercer et al. 2015), eBP_Taggart was identified from Mercer’s RNA-seq data and ENCODE RNA-seq data from 99 cell lines (GSE30567) (Taggart et al. 2017), eBP_Pineda was obtained from 17,164 RNA-seq data sets from GTEx and TCGA (Pineda and Bradley 2018), whereas eBP_Talhouarne was acquired from RNA-seq data of cytoplasmic RNA from 5 cell lines (PRJNA479418) (Talhouarne and Gall 2018). However, these previous RNA-seq studies either only described their processing steps without providing the tool, or provided a perl program with insufficient documentation. The R package LaSSO (Bitton et al. 2014), designed for lariat and BP detection from RNA-seq data, did not work in our test. The fifth dataset eBP_Briese was detected by spliceosome iCLIP experiment in 40 cell lines (E-MTAB-8182) (Briese et al. 2019).

### Identification of BP from RNA-seq data from *DBR1*-mutated patients

We obtained 15 RNA-seq data from fibroblasts of three *DBR1*-mutated patients with brainstem viral infection under five different stimulation conditions (non-stimulation (NS), IFNα, pIC, HSV1-8h, and HSV1-24h) (SRP130621), which we previously studied (Zhang et al. 2018b). *DBR1* encodes the only known lariat debranching enzyme. This RNA-seq dataset is paired-end 150-bp long, and each sample contains around 70 million reads. We mapped the fastq reads onto the human reference genome GRCh37 with STAR aligner v2.7 (Dobin et al. 2013), and outputted the unmapped reads to a new fastq file for each sample. We used Trimmomatic (Bolger et al. 2014) to remove the low-quality reads and to trim the low-quality ends, to obtain the remaining reads for BP searching. To this end, we developed our one-line command Python program (BPHunter_fastq2BP.py) embedded with BLAST+ (Camacho et al. 2009), to identify 5’ss-BP junction reads and hence BP positions (**Figure 2a, Data and Web Access**). Based on the GENCODE human reference genome GRCh37 and its gene annotation (Frankish et al. 2019), we extracted the 20-nt intronic sequences downstream of 5’ss (20-nt 5’ss library), and the 200-nt intronic sequences upstream of 3’ss (200-nt 3’ss library) for introns longer than 200 nt. For introns shorter than 200 nt, we used the entire intronic sequences in the 200-nt 3’ss library. We first aligned all BP-searching reads to the 20-nt 5’ss library, retaining the reads that had a perfect 20-nt match or only one mismatch as 5’ss-hit reads. For each 5’ss-hit read, we trimmed away the read sequence from the start of its alignment to the 20-nt 5’ss library, and inverted the remaining sequence. We then aligned the trimmed 5’ss-hit reads to the inverted 200-nt 3’ss library, and retained those reads that had at least a 20-nt alignment with at least 95% identity in the same intron, as 5’ss-3’ss-hit reads. The ends of the aligned sequence in the 200-nt 3’ss library were used to determine the genomic positions of BP. This process yielded a total of 280,899 5’ss-BP junction reads from 15 RNA-seq datasets, harboring 8,682 unique BP positions (**Table S1**).

### Consensus-guided positional adjustment of BP

Since the transesterification reaction between 5’ss and BP generates a noncanonical 2’-to-5’ linkage, and the reverse transcriptase in RNA-seq can introduce mutations (mismatches, micro-insertions/deletions) when traversing the 5’ss-BP junction (Mercer et al. 2015; Taggart et al. 2017), we anticipated that a number of eBP sites could have been mis-located in the raw dataset (**Figure 2b**). We therefore screened a window of [-3, +3] nt from each BP for consensus sequence (YTNAY) matching, and adjusted the raw BP position to its closest neighbor that perfectly matched the consensus (**Figure 2c**). If the YTNAY pattern did not match, the slightly relaxed consensus sequence (YTNA) was used to adjust the raw BP positions.

### cBP datasets

We also collected three datasets of computationally predicted BP, and we named these three datasets after their method names: cBP_BPP (Zhang et al. 2017), cBP_Branchpointer (Signal et al. 2018) and cBP_LaBranchoR (Paggi and Bejerano 2018) (**Table 1**). cBP_BPP was trained by using expectation maximization algorithm on eBP_Mercer data, and predicted BP in the 14-nt region [-21, -34] nt of 3’ss (Zhang et al. 2017). cBP_Branchpointer was trained by gradient boosting machine on eBP_Mercer data, and predicted BP in the 27-nt region [-18, -44] nt of 3’ss (Signal et al. 2018). cBP_LaBranchoR was trained by sequence-based deep-learning on eBP_Mercer and eBP_Taggart data to predict BP in the region [-1, - 70] nt of 3’ss (Paggi and Bejerano 2018).

### Prediction of BP in the region [-3, -40] nt upstream of 3’ss

As the previous BP predictions only used one or two eBP datasets for training, and overlapped the prediction in the region [-21, -34] nt of 3’ss, we supplemented them with an additional high-precision BP prediction in the region [-3, -40] nt of 3’ss that covered the sequences closer to 3’ss. We used all 194,639 consensus-guided position-adjusted eBP as positive training data (**Results**), and randomly generated 1,000,000 non-BP positions from intronic and exonic regions as negative training data. We extracted the flanking 11-nt motif [-7, +3] nt of each BP and non-BP position in the training data, based on the interaction mode between BP and snRNA (**Figure 1b**). We then vectorized the 11-nt motif into 44-bit binary code by one-hot-encoding (converting A to 0001, C to 0010, G to 0100, and T to 1000). We developed three machine learning classification models: gradient boost machine (GBM), random forest (RF), and logistic regression (LR), by using scikit-learn (Pedregosa et al. 2011) (**Figure 2d**). For each model, we first performed parameter optimization by using grid search of different combinations of key parameters, and evaluated the performance of each set of parameters by stratified-shuffled 10-fold cross-validation based on its F1 score (F1 = 2 * (precision * recall) / (precision + recall)). The parameters yielding the highest F1 score were selected as the optimal parameters (**Table S8**). As we aimed to identify the BP candidates with high precision, we performed thresholding optimization to establish the optimal probability cutoff for each model. We generated precision-recall curves (PRC) and receiver operating characteristic curves (ROC), by averaging the performance of stratified-shuffled 10-fold cross-validation. We determined the optimal threshold of each model by requiring precision ≥ 0.95 and maximizing Youden’s J statistic (J = sensitivity + specificity - 1). We therefore trained and optimized GBM-BP model, RF-BP model and LR-BP model respectively, and then combined them by majority voting for improved performance (**Table S8**). We then extracted all positions in the region [-3, -40] nt of all 3’ss, and vectorized their flanking 11-nt motif into binary code as input for BP prediction.

### Intronic data

We obtained human genome sequence and gene annotation data on the hg19/GRCh37 genome assembly from the GENCODE database (Frankish et al. 2019). By focusing on protein-coding genes and transcripts, and requiring the gene/transcript status to be known and the confidence level to be 1 or 2, we extracted a total of 43,225 alternatively spliced transcripts from 19,149 protein-coding genes. We identified multi-exon transcripts and removed introns that were shorter than 10 nucleotides, thereby obtaining 355,472 introns (200,059 unique introns) from 41,975 transcripts of 17,372 genes (**Figure S3**). We tested the genomic overlaps between these 200,059 introns, and identified 41,952 introns with alternative splicing. We also collected 672 and 752 minor introns reported by the IAOD (Moyer et al. 2020) and MIDB (Olthof et al. 2019) databases respectively, and detected 18 new minor introns by implementing the intron classification criteria proposed by MIDB (**Supplemental Data 2**). A gene ontology analysis revealed that the genes harboring minor introns were enriched in intracellular transport and ion channels.

### Mapping BP to Introns

The genomic positions of all 567,053 BP and the genomic ranges of all 200,059 introns were formatted into BED files. We then we mapped BP to introns using BEDTools (Quinlan and Hall 2010), to identify all the pairwise BP-intron associations based on their positional intersection.

### Nucleotide composition

Since U2/U12-snRNA binds to the [-5, +3] and [-7, +2] regions of BP respectively (**Figure 1b**), we defined the region of union [-7, +3] as the BP motif. We then measured the nucleotide frequency and information content (IC) of BP, 5’ss and 3’ss motifs in major and minor introns respectively, and plotted them by using SeqLogo (Bembom 2017). Information content is a measure of entropy, reflecting the degree of nucleotide conservation at each position, by heightening the highly conserved nucleotides and flattening the evenly appearing nucleotides. The IC is computed as 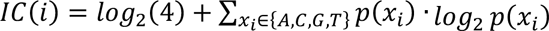, where *p(xi)* is the probability of a nucleotide *xi* in (A, C, G, T) at the position *i*. A position with a 100% conserved nucleotide has IC = 2, and a position with four equally appearing nucleotides each with 25% frequency has IC = 0.

### BP-U2/U12 snRNA binding energy

The flanking sequences of a BP undergo base-pairing with U2 or U12 snRNA, whereas the BP site itself bulges out and is not involved in the interaction with snRNA (Turunen et al. 2013; Mercer et al. 2015) (**Figure 1b**). We used the RNAcofold function from ViennaRNA package (Lorenz et al. 2011) to estimate the binding energy between BP motifs (excluding the BP sites) and U2/U12 snRNA sequences (U2: AUGAUGUG, U12: AAGGAAUGA), according to the associated intron type. RNAcofold allows the intermolecular base pairing between two RNA sequences to form static interactions, and computes the minimum free energy (MEF), which is always negative or equal to zero, to represent the binding energy (in unit: kcal/mol): a value close to zero denotes unstable binding, whereas a more negative value denotes more stable binding between the BP motif and U2/U12 snRNA.

### Motif searching in the region [-50, +20] nt surrounding BP

We searched for enriched motif patterns potentially concealed within the 50-nt upstream region and 20-nt downstream region of the BP positions separated by their nucleotides (adenine-BP, cytosine-BP, guanine-BP, and thymine-BP). We performed XSTREME analysis (Bailey et al. 2009) for motif discovery (motif width: 5-10 nt), and reported those enriched motifs having *p*-values <0.05 and appearing at >5% in each of the nucleotide-separated BP datasets. The *p*-values were computed by using the randomly shuffled nucleotides from the input sequences as the background.

### Human population variant data

We obtained human genetic variants from the gnomAD database (Karczewski et al. 2020) v3.1, which contained 76,156 WGS datasets on the hg38/GRCh38 genome assembly. We converted the variants’ genomic positions from GRCh38 to GRCh37 by using the liftover program from the UCSC Genome Browser (Kuhn et al. 2013), to allow a consistent presentation of all the genomic data in this article. By focusing on protein-coding genes, we obtained a total of 256,800,911 human variants and their total allele count (AC) and general minor allele frequencies (MAF). We categorized variants into singleton (AC = 1), rare (MAF < 1e-3), low-frequency (MAF: 1e-3 ∼ 0.05), and common (MAF > 0.05).

### Cross-species conservation scores

We obtained the pre-computed genome-wide cross-species conservation scores (GERP and PhyloP-46way) from the UCSC Genome Browser (Kuhn et al. 2013). GERP (Genomic Evolutionary Rate Profiling) computes position-specific scores of evolutionary constraint using maximum likelihood evolutionary rate estimation by aligning 35 mammals (Cooper et al. 2005). PhyloP-46way (Phylogenetic P-values) measures the evolutionary base-wise conservation based on the alignment of 46 vertebrates (Pollard et al. 2010). Both scores indicate the strength of purifying selection of a given genomic position: a positive value denotes that the genomic position is likely to be evolutionarily conserved across species, whereas a negative value indicates that the genomic position is probably evolving neutrally.

### Mutation deleteriousness, mis-splicing prediction scores, and splice site strength

We used CADD v1.6 (Rentzsch et al. 2019), which predicts the deleteriousness of variants by taking account of an array of nucleotide sequence information and variant annotations (including conservation, amino acid change, epigenetic modification, human population variation, splicing, etc.). We extracted CADD PHRED-scaled scores (ranging from 0 to 99): the larger the score, the higher possibility of deleteriousness. It precomputed and ranked all possible variants in the human genome, and then assigned a score of 10 to the top 10% of predicted deleterious variants, a score of 20 to the top 1% of variants, and a score of 30 to the top 0.1% of variants, etc. Usually, a high-cutoff 20 or a moderate-cutoff 10 were used for large-scale variant filtration for deleterious candidate mutations (Zhang et al. 2018a; Zhang et al. 2021). We were also aware of some mis-splicing prediction scores (Xiong et al. 2015; Cheng et al. 2019; Jagadeesh et al. 2019; Jaganathan et al. 2019), and in this study we recruited SpliceAI (Jaganathan et al. 2019) and MMSplice (Cheng et al. 2019) to evaluate the BP mutations. SpliceAI predicts splice junctions from RNA sequence, by means of a deep neural network model (Jaganathan et al. 2019). It provides a score (ranging from 0 to 1) for disrupting the acceptor site: the higher the score, the higher the probability of altering the acceptor site. SpliceAI suggested a high-cutoff 0.8 for high-precision, and also recommended a moderate- cutoff 0.5. MMSplice predicts the effect of variants on splicing, by means of a neural network-based modular modelling on different components of splicing. It computes a score (unspecified range) for acceptor site inclusion (positive score) or exclusion (negative score). MMSplice suggested a cutoff of -2 to be considered as evidence for acceptor site disruption (Cheng et al. 2019). In studying splice site strength, we used SeqTailor (Zhang et al. 2019) to extract the wild-type (wt) and mutated (mt) 23-nt DNA sequences surrounding the mutations of interest, and then used MaxEntScan (Yeo and Burge 2004) to estimate the splice site strength in wt and mt sequences respectively.

### A cohort of patients with critical COVID-19

In this study, we used the WES data from a cohort of 1,035 patients with critical COVID-19, which were recruited through an international consortium - The COVID Human Genetic Effort (Casanova et al. 2020). All human subjects in this study were approved by the appropriate institutional review board.

### Exon trapping assay

DNA segments encompassing *STAT2* exon 5 and 6 region (chr12:56749479 to chr12:56748872 region, GRCh37 reference) were amplified from genomic DNA extracted from PBMCs of a healthy control and were cloned into a pSPL3 vector, using the *EcoR*I and *BamH*I sites. c.472-24 A>T of *STAT2* (an intronic variant located in intron 5 and predicted to alter a branchpoint) was generated by site-directed mutagenesis. Plasmids containing wild-type and mutant *STAT2* exon 5 and 6 region were then used to transfect COS-7 cells. After 24 hours, total RNA was extracted and reverse transcribed. cDNA products were amplified using flanking HIV-TAT sequences of the pSPL3 vector, and ligated into the pCRTM4-TOPO® vector (Invitrogen). StellarTM cells (Takara) were transformed with the resulting plasmids. Colony PCR and sequencing using primers located in the flanking HIV-TAT sequences of the pSPL3 were performed to assess the splicing products transcribed by the wt and mutant alleles.

### TOPO-TA cloning and RT-qPCR

Total RNA was extracted from a whole blood sample from the patient and a healthy control using Tempus Blood RNA Tube and Tempus Spin RNA Isolation Kit (Applied Biosystems), and reverse transcribed into cDNA using SuperScript III (Invitrogen). For TOPO-TA cloning, specific primers located in exon 3 (forward primer, CATGCTATTCTTCCACTTCTTG) and exon 7-8 boundaries (reverse primer, GGCATCCAGCACCTCCTTTC) were used to amplify *STAT2* cDNA by PCR. PCR products were then purified and ligated into a pCRTM4-TOPO® vector (Invitrogen). StellarTM cells (Takara) were transformed with the resulting plasmids. Colony PCR and sequencing using the primers used to amplify *STAT2* cDNA were performed to assess the splicing products generated from the wt and mutated alleles. For RT-qPCR, *STAT2* mRNAs were quantified using probes Hs01013129_g1 (exons 5-6) and Hs01013130_g1 (exons 6-7; Thermo Fischer Scientific), with the Taqman Gene Expression Assay (Applied Biosystems), and normalized to the expression level of human β-glucuronidase. Results were expressed using the ΔΔCt method, as described by the manufacturer, and the amount of *STAT2* canonical transcript (ENST00000314128.9) was estimated based on the TOPO-TA data using the following formula: ΔΔCt x percentage of canonical transcript/percentage of transcripts with canonical exon 5-6 junction for probe Hs01013129_g1 or ΔΔCt x percentage of canonical transcripts/percentage of transcripts with canonical exon 6-7 junction for probe Hs01013130_g1.

### A cohort of lymphoma patients

We studied the somatic mutations from a cohort of 53 diffuse large B-cell lymphoma patients, whose paired WGS and RNA-seq data from the tumor tissues were also available (Georgiou et al. 2016; Ren et al. 2018; Ye et al. 2021). We focused on a set of 212 genes that are frequently mutated in B-cell lymphomas or known to be important for B-cell lymphomagenesis (Ren et al. 2018). All human subjects in this study were approved by the appropriate institutional review board.

### COSMIC database

We collated the somatic mutations documented in the COSMIC database (Tate et al. 2019) v94, which have been detected in cancer patients from a variety of different sources. We also identified four gene sets of interest, which were related with cancer formation and progression: 123 tumor suppressor genes, 161 apoptosis genes, 150 DNA repair genes, and 714 cell cycle genes, based on COSMIC (Tate et al. 2019) and MSigDB (Liberzon et al. 2015) databases. We used the following criteria to retain the high-confidence candidate BP mutations: (1) canonical transcripts; (2) deletions and SNVs that remove or disrupt the entire BP motifs or the BP/BP-2 positions; (3) in introns harboring single or two BP; (4) no or very rare (MAF < 1e-3) population variations.

## DATA AND WEB ACCESS

BPHunter data and software are publicly accessible from http://hgidsoft.rockefeller.edu/BPHunter and https://github.com/casanova-lab/BPHunter. It supports both hg19/GRCh37 and hg38/GRCh38 human reference genomes. Given a VCF file of variants with its first five columns as mandatory fields (CHROM, POS, ID, REF, ALT) as input, BPHunter outputs a text file of the variants that may disrupt BP with the following informative annotations: variant type (snv / *x* nt-deletion / *x* nt-insertion), gene symbol, transcript, intron number, intron length, intron type (major / minor), BP name, BP ranking among the total number of BP in this intron, BP positions affected ([-2, 0] of BP), distance from BP to 3’ss, population variant MAF of BP, GERP and PhyloP conservation scores of BP, BP-snRNA binding energy, level of consensus (1:YTNAY / 2:YTNA / 3:TNA / 4:YNA), evidence score (EVI: a two-digit score: xy, where x = number of eBP sources, y = number of cBP sources), and list of supporting sources. The negative BP reporting system in BPHunter webserver requires the submitters to provide their personal contact details (name, Email and institution), variant information (genome assembly, chromosome, position, reference allele, mutant allele and gene symbol), BP name (as provided by BPHunter), cell type, assay type and experimental evidence following the given template (**Supplemental Data 7**).

## ACKNOWLEDGEMENT

We thank Y. Nemirovskaya, M. Woollett and L. Lorenzo for administrative support, A. Cobat and E. Jouanguy for assistance, and Z. Yang and Y. Nemirovskaya for artwork design. The Laboratory of Human Genetics of Infectious Diseases is supported by the Howard Hughes Medical Institute, the Rockefeller University, the St. Giles Foundation, the National Institutes of Health (NIH) (R01AI088364), the National Center for Advancing Translational Sciences (NCATS), NIH Clinical and Translational Science Award (CTSA) program (UL1 TR001866), the Yale Center for Mendelian Genomics and the GSP Coordinating Center funded by the National Human Genome Research Institute (NHGRI) (UM1HG006504 and U24HG008956), the French National Research Agency (ANR) under the “Investments for the Future” program (ANR-10-IAHU-01), ANR grants (ANR-14-CE14-0008-01, ANR-18-CE15-0020-02 and ANR-20-CO11-0001), the Integrative Biology of Emerging Infectious Diseases Laboratory of Excellence (ANR- 10-LABX-62-IBEID), the French Foundation for Medical Research (FRM) (EQU201903007798), a Fast Grant from Emergent Ventures, Mercatus Center at George Mason University, the Fisher Center for Alzheimer’s Research Foundation, the Meyer Foundation, the JPB Foundation, the FRM and ANR GENCOVID project (ANR-20-COVI-0003), ANRS Nord-Sud (ANRS-COV05), the Square Foundation, Grandir-Fonds de solidarité pour l’enfance, the SCOR Corporate Foundation for Science, The Foundation du Souffle, *Institut National de la Santé et de la Recherche Médicale* (INSERM) and the University of Paris. We also thank the Swedish Cancer Society (Cancerfonden), the Swedish Research Council, Radiumhemmets, the Center for Innovative Medicine (CIMED) and the Knut and Alice Wallenberg Foundation.

